# Rapid binding to protofilament edge sites facilitates tip tracking of EB1 at growing microtubule plus-ends

**DOI:** 10.1101/2022.06.07.495114

**Authors:** Samuel J. Gonzalez, Rebecca R. Goldblum, Katherine T. Vu, Rachel Shoemaker, Taylor Reid, Mark McClellan, Melissa K. Gardner

**Author notes:** Correspondence: Melissa K Gardner.

## Abstract

EB1 is a key cellular protein that delivers critical regulatory molecules throughout the cell via the tip-tracking of growing microtubule plus-ends. Thus, it is important to understand the mechanism for how EB1 efficiently tracks growing microtubule plus-ends. It is widely accepted that EB1 binds with higher affinity to GTP-tubulin subunits at the growing microtubule tip, relative to GDP-tubulin along the microtubule length. However, it is unclear whether this difference in affinity alone is sufficient to explain the tip-tracking of EB1 at growing microtubule tips. Previously, we found that EB1 binds to exposed microtubule protofilament-edge sites at a ∼70-fold faster rate than to closed-lattice sites, due to diffusional steric hindrance to binding. Thus, we asked whether rapid protofilament-edge binding could contribute to efficient EB1 tip tracking. A computational simulation with differential EB1 on-rates based on closed-lattice or protofilament-edge binding, and with EB1 off-rates that were dependent on tubulin hydrolysis state, recapitulated experimental EB1 tip tracking. To test this model, we used cell-free biophysical assays, as well as live-cell imaging, in combination with the chemotherapy drug Eribulin. We found that Eribulin blocked EB1 protofilament-edge binding, which led to a dramatic decrease in EB1 tip tracking on dynamic microtubules. We conclude that rapid EB1 binding to microtubule protofilament-edge sites increases the efficiency of EB1 tip tracking at the growing microtubule plus end.

## Introduction

Microtubules are important cellular filaments composed of αβ tubulin heterodimers that are stacked end-to-end to form structures known as protofilaments. These protofilaments associate laterally to form the hollow-tube structure of the microtubule (Mitchison & Kirschner, 1984). The α/β polarity of the tubulin dimer induces microtubule polarity, such that the microtubule end with β-tubulin exposed forms the fast-growing, dynamic “plus-end” of the microtubule (Desai & Mitchison, 1997; Mitchison & Kirschner, 1984). In solution, the β-tubulin subunit binds to a GTP nucleotide, which then hydrolyzes to GDP after incorporation into the microtubule lattice. This delayed hydrolysis leads to a high concentration of GTP tubulin at growing microtubule plus-ends, commonly referred to as the “GTP-cap” (Desai & Mitchison, 1997). The presence of the GTP-cap creates a distinct region that is present exclusively at growing microtubule plus-ends (Maurer et al., 2011, 2014; Zanic et al., 2009).

The localization of proteins along different regions of the microtubule is central to the role of microtubules in cell migration, intracellular transport, and cell division (Barlan & Gelfand, 2017; Burute & Kapitein, 2019; Etienne-Manneville, 2013; Farmer et al., 2021; Forth & Kapoor, 2017; Garcin & Straube, 2019; Goodson & Jonasson, 2018; Kaverina & Straube, 2011; Martin & Akhmanova, 2018; Maurer et al., 2012, 2014; Siddiqui & Straube, 2017). EB1 is a key cellular protein that autonomously localizes to the growing ends of microtubules (“tip tracks”) and recruits other important proteins that have little or no affinity to growing microtubule ends (Bieling et al., 2007; Dixit et al., 2009; Morrison et al., 1998; Mustyatsa et al., 2017). It has been shown that improper localization of EB1 at growing microtubule plus-ends can lead to disruptions in both cell division and cell migration (Dong et al., 2010; Honoré et al., 2008; Mustyatsa et al., 2017; Rogers et al., 2002).

EB1 binds a small pocket within the microtubule lattice that is created by four tubulin dimers. Importantly, it has been shown that EB1 binds with a higher affinity to GTP-tubulin subunits than for GDP tubulin subunits. (Maurer et al., 2011, 2012, 2014; Zanic et al., 2009; Zhang et al., 2015). This difference in affinity has been shown to increase the enrichment of EB1 within the GTP-cap at growing microtubule plus-ends.

Recent work has demonstrated that EB1 can also bind to a partial binding pocket composed of 2-3 tubulin subunits, either at the distal end of the growing microtubule, or at lattice openings within the microtubule (Reid et al., 2019). We define these exposed, partial binding pockets as “protofilament-edge” sites. Importantly, we recently reported that the arrival rate of EB1 to 2- tubulin protofilament-edge sites was ∼70-fold faster than to complete 4-tubulin pockets, due to a reduced diffusional steric hindrance to binding (Reid et al., 2019). Because protofilament- edge sites are present at growing microtubule plus-ends (Atherton et al., 2018; Gudimchuk et al., 2020; Guesdon et al., 2016), we hypothesized that this large difference in EB1 arrival rates could have important repercussions for the efficiency of EB1 tip tracking at growing microtubule plus-ends. Specifically, we predicted that the rapid binding of EB1 to protofilament-edge sites at the growing microtubule plus-end could increase the efficiency of EB1 plus-end tip tracking.

In this work, we generated a stochastic simulation that incorporated the assembly and hydrolysis of tubulin subunits, as well as the binding and unbinding of EB1 molecules. Importantly, in our simulation, EB1 rapidly bound to protofilament-edge sites, and bound more slowly to closed-lattice sites. In addition, consistent with previous affinity measurements, the off-rate of EB1 bound to closed pocket GTP-tubulin sites was low, with higher EB1 off-rates from GDP-tubulin pocket sites. This simulation predicted that rapid binding to protofilament- edge sites was required for EB1 to efficiently tip-track growing microtubule plus-ends. To test this prediction, we used cell-free biophysical assays, as well as live cell imaging, in combination with the chemotherapy drug Eribulin. Consistent with previous studies, we found that Eribulin binds to protofilament-edge sites on microtubules (Doodhi et al., 2016; Smith et al., 2010).

Further, Eribulin blocked EB1 protofilament-edge binding, which led to a dramatic decrease in EB1 tip tracking on dynamic microtubules, both in cell-free experiments and in cells. Together, this work demonstrates that protofilament-edge binding, together with a differential EB1 affinity for GTP and GDP tubulin, facilitates efficient EB1 tip tracking.

## Results

### Stochastic simulations with rapid binding at protofilament-edge sites can recapitulate EB1 tip tracking

In previous work, we found that the arrival rate of EB1 to exposed protofilament-edge sites was ∼70-fold faster than to closed four-tubulin pockets, due to a diffusional steric hindrance to binding (Reid et al., 2019). To ask whether rapid EB1 protofilament-edge binding could contribute to EB1 tip tracking, we created a stochastic simulation in which there was an increased on-rate of EB1 to protofilament-edge sites relative to closed-lattice sites. This simulation combined the assembly of individual tubulin subunits, together with EB1 arrivals, binding, and departures from the microtubule.

The tubulin assembly portion of the model was built on earlier work, in which individual tubulin subunits were allowed to arrive and depart from the growing microtubule plus-end, and lattice-incorporated tubulin subunits were stochastically hydrolyzed at a constant rate (C. E. Coombes et al., 2013; Margolin et al., 2011, 2012; VanBuren et al., 2005) (see Methods).

In addition to tubulin subunit assembly, individual EB1 molecules were allowed to bind and unbind from their binding pockets at any position on the growing microtubule (Fig. 1A).

**Figure 1.**
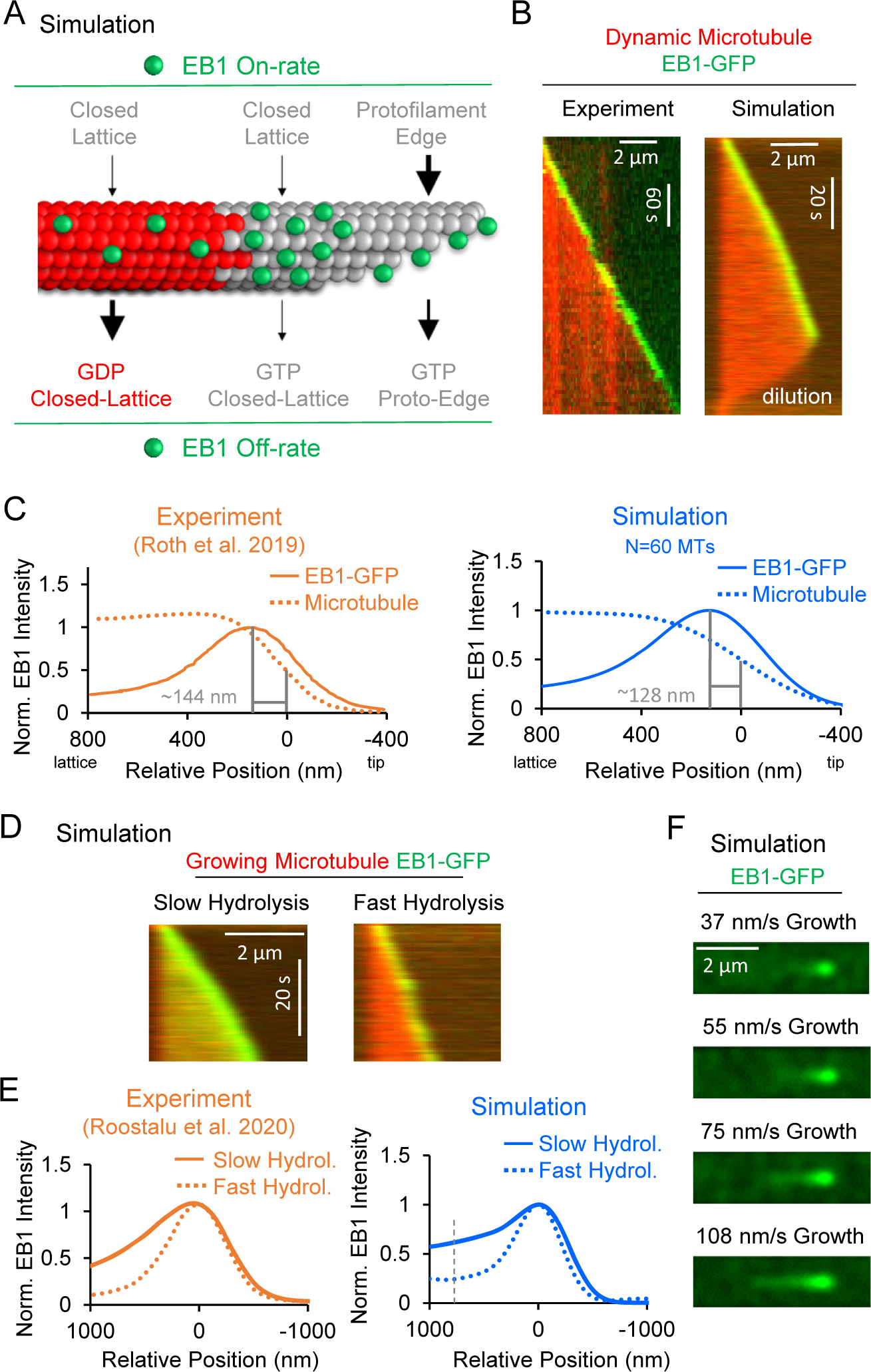
Development and validation of a stochastic simulation for EB1 tip tracking. (A) Rules for a molecular-scale stochastic simulation that incorporates both tubulin subunit assembly and EB1 arrivals and departures from the growing microtubule (See Methods). In the simulation, the EB1 protofilament-edge on-rate (top-right) is substantially higher than the EB1 closed- lattice on-rate (top-left and top-center). The EB1 off-rate is faster for GDP-tubulin binding pockets (bottom-left) than for GTP-tubulin binding pockets (bottom-center). In addition, the EB1 protofilament-edge off-rate is faster than the EB1 closed-lattice off-rate (bottom-right). (B) Left: Experimental EB1 tip tracking. EB1-GFP is strongly localized to growing microtubule ends. Right: Simulated EB1 tip tracking at growing microtubule ends. (C) Left: Line scans of EB1 intensity from experimentally reported data (Roth et al., 2019) (orange). Right: Line scans of EB1 intensity from the simulation (blue, see Methods). (D) Left: Simulated EB1 tip tracking with slow GTP-tubulin hydrolysis rate (0.1 s^-1^) Right: Simulated EB1 tip tracking with a fast GTP-tubulin hydrolysis rate (2 s^-1^) (E) Left: Experimental line scan quantification of Mal3-GFP intensity along the length of the microtubule at two different hydrolysis rates (orange, (Roostalu et al., 2020)). Right: Simulated data line scan quantification of EB1-GFP intensity along the length of the microtubule at two different hydrolysis rates (blue). (F) Increasing growth rate in the simulation increases the EB1 comet length, similar to reports in the literature (Maurer et al., 2014).

However, the EB1 on-rates and off-rates depended on the individual binding pocket chemistry and configuration. Specifically, the on-rates for individual EB1 molecules depended on the structure of the binding pocket, where EB1 arrivals to protofilament-edge sites were substantially faster than to closed-lattice sites, regardless of hydrolysis state (Fig. 1A, top, see Methods) (Reid et al., 2019). In contrast, the off-rate of bound EB1 molecules depended on the hydrolysis state of the EB1 binding site. Here, the tubulin subunits towards the minus end of the EB1 binding site dictated the “hydrolysis state” of the EB1 binding site. If 1-2 of these tubulin subunits were hydrolyzed to GDP, the binding site was considered to be a “GDP” binding site, leading to an increased EB1 off-rate (see Methods). We note that, because the protofilament-edge binding sites are incomplete (e.g., lacking the entire 4-tubulin pocket), the EB1 off-rate from GTP-tubulin protofilament-edge sites was ∼10-fold higher than for GTP-tubulin closed-lattice sites.

We first asked whether EB1 “tip tracked” growing microtubule plus-ends in the simulation, similar to experimental observations (Fig. 1B, left). Qualitatively, our simulated EB1 behaved similarly to experiments – strongly targeting growing microtubule plus-ends, while detaching from shortening ends (Fig. 1B, right, Movie S1). To quantitatively confirm that the simulated EB1 tip tracking was similar to experimental results, we next compared the “peak EB1 intensity position” from our simulation data, to results reported in the literature. The peak EB1 intensity position refers to the location of the highest EB1 intensity relative to the growing microtubule plus-end (Maurer et al., 2012, 2014; Nakamura et al., 2012; Roth et al., 2019). At a microtubule growth rate of 10-30 nm/s, the peak EB1 position has been reported to be ∼144 nm from the microtubule end (Fig. 1C, left, orange) (Roth et al., 2019). To quantify the peak EB1 position in the simulation, line scans of simulated EB1 comets were obtained and averaged over 60 simulation runs. We found that the simulation predicted a peak EB1 intensity position of ∼128 nm, similar to experimental observations (Fig. 1C, right, blue; growth rate 35 nm/s).

To ensure that the configuration of the microtubule plus-end was similar between experiment and simulation, we compared the fitted tip standard deviation in simulated microtubule images to previously reported experimental values. We found that the average tip standard deviation of our simulated microtubules was ∼155 nm, which was similar to previously reported in vitro values of ≤145 nm (Maurer et al., 2014) (Fig. S1A).

It has been shown that a suppressed GTP-tubulin hydrolysis rate increases the EB1-GFP intensity at the microtubule growing end, likely due to an increased GTP-cap size (Roostalu et al., 2020). Thus, to ask whether the simulation could recapitulate this phenomenon, we ran simulations with both slow and fast GTP-tubulin hydrolysis rates. We found that a slower hydrolysis rate led to an increased concentration of EB1 at the growing microtubule plus-end, while a faster hydrolysis rate led to a reduced concentration of EB1 at the growing microtubule end, similar to previous reports (Fig. 1D, Movie S2) (Roostalu et al., 2020). By quantifying the localization of EB1 at growing microtubule plus-ends in these simulations, we observed a ∼2.5- fold increase in EB1 binding along the lattice of simulated microtubules with a slow hydrolysis rate, relative to a fast hydrolysis rate (Fig. 1E, right, blue; calculated at position 768 nm, grey dashed line), similar to previously reported experimental results (Fig. 1E, left, orange; (Roostalu et al., 2020)). This result demonstrates that EB1 tip tracking in the simulation depends on the tubulin hydrolysis rate, similar to previous experimental results (Roostalu et al., 2020).

Finally, it has been widely reported that an increased microtubule growth rate leads to a longer EB1 “comet” (Farmer et al., 2021; Maurer et al., 2014; Reid et al., 2019). Thus, we ran simulations with increasing microtubule growth rates, keeping the GTP-tubulin hydrolysis rate constant. Similar to experimental reports, we found that, as the microtubule growth rate was increased in the simulation, the comet length was increased (Fig. 1E) (Farmer et al., 2021; Maurer et al., 2014; Reid et al., 2019).

In summary, multiple characteristics of EB1 tip tracking from the literature were recapitulated in simulations that included rapid EB1 protofilament-edge binding. To evaluate the robustness of EB1 tip tracking in the simulation, we performed parameter testing with respect to the EB1 on-rates and off-rates (Fig. S1B-L, Fig. S2). We found that even the most sensitive parameters had at least an 8-fold range of acceptable values in which EB1 was clearly localized exclusively at the growing microtubule end (Fig. S1B-L, Fig. S2).

### Simulations require protofilament-edge binding to qualitatively recapitulate tip tracking

We next asked whether, if EB1 bound exclusively to the canonical closed-lattice sites on the microtubule, the simulation could recapitulate experimentally observed EB1 tip tracking. Thus, we set the EB1 protofilament-edge on-rate to zero, and then slowly increased the EB1 closed- lattice on-rate, while leaving all EB1 off-rates constant and at baseline values (Fig. 2A; Movie S3; See Methods). We found that, while a higher EB1 closed-lattice on-rate alone led to EB1 accumulation at the growing microtubule end, it also led to substantial EB1 accumulation along the length of the microtubule (Fig. 2A), thus reducing the specificity of EB1 localization to the growing microtubule end. We note that, by substantially increasing the off-rate of EB1 from the GDP-tubulin closed-lattice, we were able to partially suppress this phenomenon (Fig. S3A).

**Figure 2.**
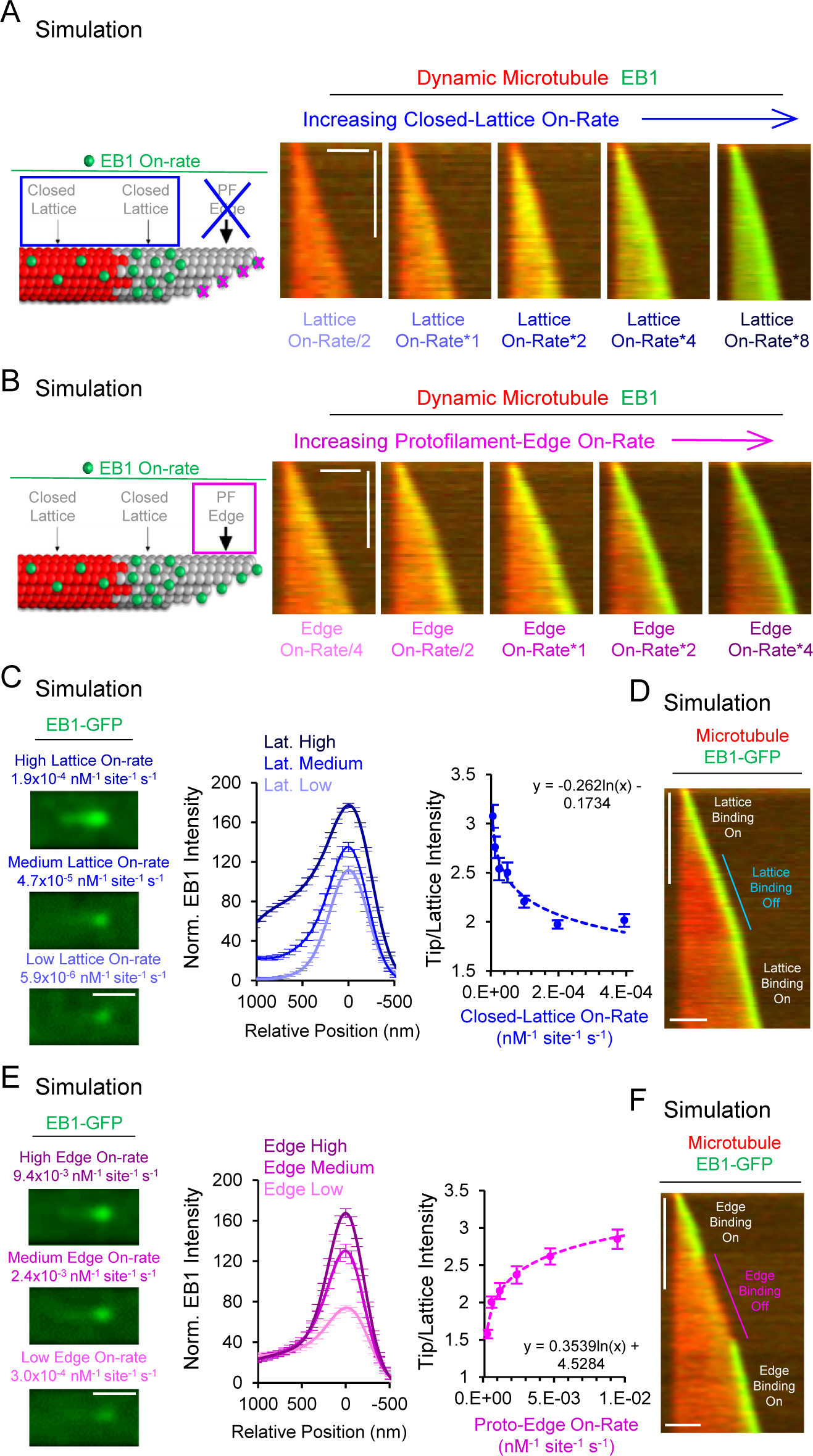
Simulations predict that protofilament-edge binding is required for robust EB1 tip tracking. (A) Left: Simulations were performed in which the protofilament-edge on-rate was set to zero, and the closed-lattice on-rates were varied. Right: Simulated kymographs in which the protofilament-edge binding on-rate was set to zero, and the on-rate at closed-lattice sites was gradually increased (scale bars: 2 µm horizontal; 10 s vertical). (B) Left: Simulations were performed in which the closed-lattice on-rate remained constant and at the low baseline value, and the protofilament-edge on-rates were varied. Right: Simulated kymographs in which the closed-lattice on-rates remained constant, and the protofilament-edge on-rates were gradually increased (scale bars: 2 µm horizontal; 10 s vertical). (C) Left: Simulated images of EB1-GFP tip tracking over a range of closed-lattice on-rates (scale bar: 1 µm). Center: Line scans from simulated images of EB1-GFP intensity for a range of closed-lattice on-rates. Right: Normalized tip/lattice EB1-GFP intensity vs closed-lattice on-rates in the simulation. Localization to the microtubule tip decreases with increasing closed-lattice on-rates. (D) Simulated kymograph in which the closed-lattice on-rate is set to zero partway through the simulation, and later returned to its baseline value. 2 μm scale bar and 10 s time scale bar. (E) Left: Simulated images of EB1-GFP tip tracking over a range of protofilament-edge on-rates (scale bar: 1 µm). Center: Line scans from simulated images of EB1-GFP intensity for a range of protofilament-edge on-rates. Right: Normalized tip/lattice EB1-GFP intensity vs protofilament-edge on-rates in the simulation. Localization to the microtubule tip increases with increasing protofilament-edge on-rates. (F) Simulated kymograph in which the protofilament-edge on-rate is set to zero partway through the simulation and later returned to its baseline value. 2 μm scale bar and 10 s time scale bar.

However, in this case, the location of the peak EB1 intensity relative to the microtubule tip diverged substantially from published experimental observations (Fig. S3A). Specifically, in simulations with a high closed-lattice EB1 on-rate (without EB1 protofilament-edge binding), the peak EB1 position ranged from ∼400-700 nm distal of the growing microtubule end, as compared to experimental measurements of ∼144 nm (Fig. S3B vs Fig. 1C) (Maurer et al., 2014; Roth et al., 2019). Thus, the simulation predicts that closed-lattice binding alone cannot recapitulate experimental observations of EB1 tip tracking.

We next explored the effect of increasing the EB1 protofilament-edge on-rates. Thus, we set the EB1 closed-lattice on-rate to its low baseline value, and then slowly increased the EB1 protofilament-edge on-rate, while leaving all EB1 off-rates constant and at baseline values (Fig. 2B, Movie S4; See Methods). We found that an increasingly intense, simulated EB1-GFP puncta appeared at the growing microtubule end as the protofilament-edge on-rate was increased (Fig. 2B, Movie S4). Importantly, the peak EB1 position ranged from 141-217 nm distal to the growing microtubule end over a 16-fold range of protofilament-edge on-rates, which was similar to the experimental reports of ∼144 nm (Fig. S3B vs Fig. 1C).

### Specificity of EB1 targeting to growing microtubule tips is reduced with higher EB1 closed-lattice binding site on-rates

To quantitatively dissect the relative role of closed-lattice binding on EB1 localization to growing microtubules, we ran simulations over a range of EB1 closed-lattice on-rates, while keeping all other EB1 on-rates and off-rates constant and set at their base simulation values, including rapid EB1 protofilament-edge binding (see Methods). We found that a low EB1 closed-lattice on-rate led to a clear EB1 puncta at the tip of the microtubule (Figure 2C, left-bottom). Here, EB1 accumulation is dominated by protofilament-edge binding. However, increasing the EB1 closed-lattice on-rate by 32-fold led to a ∼1.6-fold increase in EB1 intensity at the microtubule tip, but, importantly, also led to a ∼25-fold increase in EB1 lattice intensity along the length of the microtubule (Figure 2C, center), even in the presence of EB1 protofilament-edge binding. By plotting the ratio of EB1 microtubule tip intensity to EB1 lattice intensity (see Methods), we found that the relative EB1 intensity at the microtubule tip decreased with increasing EB1 closed-lattice on-rates (Fig. 2C, right). Thus, the efficiency of simulated EB1 tip tracking was reduced with higher EB1 binding to closed-lattice sites, due to increased EB1 accumulation along the length of the microtubule rather than only at the growing tip.

We then performed a simulation where the closed-lattice on rate was set to zero partially through the simulation, to observe in real-time the importance of the closed-lattice binding on EB1 tip tracking. We found that EB1 tip tracking appeared unaltered when closed-lattice binding was shut off during the dynamic microtubule simulation (Fig. 2D, blue; Movie S5).

### Simulations with increasing EB1 protofilament-edge on-rates lead to EB1 accumulation exclusively at the growing microtubule plus-end

Next, to quantitatively assess the role of protofilament-edge binding on EB1 localization, we ran simulations over a range of protofilament-edge on-rates, while keeping all other EB1 on-rates and off-rates constant and set at their base simulation values, including the closed-lattice on-rate (see Methods). We found that, by substantially decreasing the protofilament-edge on-rate, the intensity of EB1 at the growing microtubule tip was dimmed (Figure 2E, left-bottom). Upon increasing the protofilament-edge on-rate, the intensity of EB1 at the growing tip was clearly increased, without an increase in EB1 intensity along the length of the microtubule (Fig. 2E, left-top). Here, a 32-fold increase in protofilament-edge on-rate led to a ∼2.2-fold increase in EB1 intensity at the tip of the microtubule, and, importantly, no change in the EB1 intensity along the closed lattice of the microtubule (Fig. 2E, center). By plotting the ratio of EB1 tip intensity over EB1 lattice intensity (see Methods), we found that the relative EB1 tip intensity was increased with increasing EB1 protofilament-edge on-rates (Fig. 2E, right). Thus, the efficiency of simulated EB1 tip tracking was enhanced by higher EB1 on-rates to incomplete, protofilament-edge binding sites.

Finally, we performed a simulation where the protofilament-edge on rate was set to zero partially through a simulation. We found that EB1 tip tracking was rapidly diminished when protofilament-edge binding was shut off during a dynamic microtubule simulation, and returned very quickly when the EB1 protofilament-edge on-rate was reset to its baseline value (Fig. 2F, magenta; Movie S6).

### Chemotherapy drug Eribulin binds to protofilament edge sites on microtubules

Because the simulation predicted that protofilament-edge binding is integral to EB1 tip tracking at growing microtubule tips, we reasoned that EB1 tip tracking would be disrupted by blocking protofilament-edge sites (Fig. 3A, magenta). It has been previously shown that the chemotherapy drug Eribulin binds to protofilament-edges, and partially overlaps with the EB1 binding site on tubulin (Doodhi et al., 2016; Jordan et al., 2005; Smith et al., 2010). Specifically, EB1 binds to the H3 loop of β-tubulin, and the loss of this interaction abrogates the ability of the yeast EB1 homolog Mal3 to tip track microtubules (Maurer et al., 2012). Importantly, Eribulin also binds in the H3 loop region along β-tubulin (Doodhi et al., 2016; Maurer et al., 2012). Therefore, Eribulin could bind to protofilament-edge sites, and thus suppress EB1 binding in these locations (Doodhi et al., 2016; Maurer et al., 2012). Thus, we hypothesized that EB1 binding to protofilament-edge sites would be suppressed in the presence of Eribulin, which would in turn disrupt proper EB1 tip tracking (Fig. 3A).

**Figure 3.**
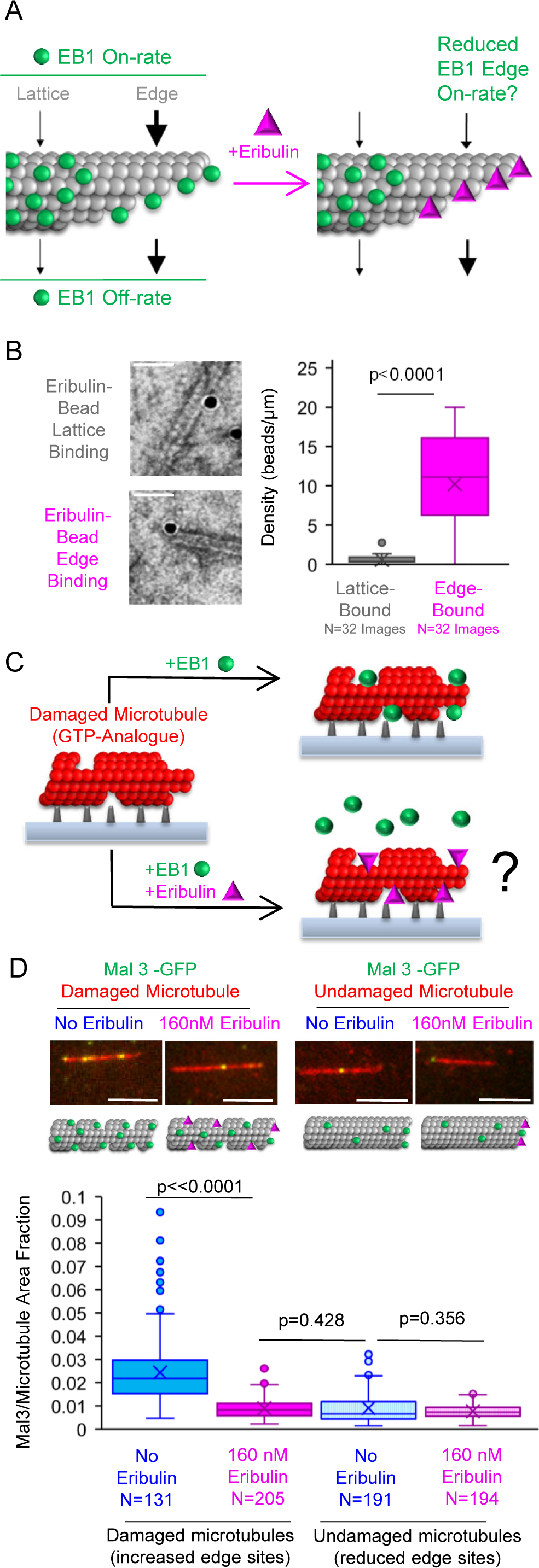
Chemotherapy drug Eribulin blocks EB1 binding to protofilament-edge sites. (A) While previous work has demonstrated that EB1 is able to bind protofilament-edge sites (left), we hypothesized that the chemotherapy drug Eribulin could bind protofilament-edge sites, and thus block EB1 binding to these sites (right). (B) Left: Electron microscopy images of Eribulin functionalized to 20 nM gold beads, visualized on taxol-stabilized microtubules. Examples of Eribulin binding along the microtubule closed-lattice (top) or at microtubule protofilament-edge sites (bottom). Right: Quantification of Eribulin-bound gold bead density on the microtubule closed-lattice (grey) or on protofilament-edges (magenta). Values were normalized to the length of each microtubule component (lattice or edge) in each image (Mann-Whitney U Test, p<<0.0001). (C) To test whether Eribulin blocks EB1 binding to protofilament-edge sites, we first generated stabilized GTP-analogue (GMPCPP) microtubules that were damaged with CaCl_2_ treatment, thus creating protofilament-edge sites along the length of the microtubule (left). Then, the microtubules were incubated with either EB1-GFP or Mal3-GFP in the presence and absence of 160 nM Eribulin (right). Loss of EB1 binding to the microtubule in the presence of Eribulin suggests that Eribulin blocks EB1 from binding to protofilament-edge sites (right, bottom). (D) Top: Typical images of Mal3-GFP binding to damaged (left) and undamaged (right) microtubules, in the absence (blue) or presence (magenta) of 160 nM Eribulin (scale bars: 5 µm). Middle: Cartoons depicting relative binding of EB1 and Eribulin in each experiment. Bottom: Quantification of the fraction of microtubule area bound by Mal3-GFP for damaged and undamaged microtubules in the absence or presence of Eribulin (ANOVA, p<<0.0001; Tukey’s post-hoc for individual comparisons; sample size indicates number of images)

To investigate this hypothesis, we first performed experiments to verify that Eribulin binds to protofilament-edge sites, as has been previously shown (Doodhi et al., 2016; Jordan et al., 2005; Smith et al., 2010). Thus, we conjugated Eribulin to 20 nm gold beads (see Methods), and then performed negative stain Transmission Electron Microscopy (TEM) to identify the location of Eribulin-bead binding on Taxol-stabilized microtubules (Fig. 3B, left). We found that the density of Eribulin-beads bound to putative protofilament-edge sites was ∼20-fold higher than for Eribulin-beads bound to closed-lattice sites (Fig. 3B, right; p<<0.0001, t-test, beads per unit length of microtubule), confirming previous observations (Doodhi et al., 2016).

### Chemotherapy drug Eribulin suppresses EB1 binding to protofilament-edge sites on microtubules

Since Eribulin binds to protofilament-edge sites, we next asked whether Eribulin could block EB1 binding at these same sites. To do this, we first generated stabilized GTP-analogue microtubules that were damaged, such that portions of the microtubule contained openings and defects. Damaging the microtubules in this way leads to an increased density of protofilament-edge sites along the microtubule length (Fig. 3C, left) (C. Coombes et al., 2016; Gupta et al., 2013; Reid et al., 2017). Opened, damaged microtubules can be generated by briefly exposing GTP-analogue (GMPCPP) microtubules to CaCl_2_ (C. Coombes et al., 2016; Gupta et al., 2013; Reid et al., 2017). Therefore, coverslip adhered, rhodamine-labeled GMPCPP microtubules were briefly incubated in 40 uM CaCl_2_, followed by a wash. Then, the damaged microtubules were incubated with either EB1-GFP or its yeast equivalent Mal3-GFP, in the absence or presence of Eribulin (Fig. 3C). Total Internal Reflection Fluorescence (TIRF) microscopy was then used to visualize the microtubules, and protein binding to the microtubules was assessed (Fig. 3C, right). Qualitatively, we observed a dramatic reduction in Mal3-GFP binding to the damaged microtubules in the presence of Eribulin, as compared to the no-Eribulin controls (Fig. 3D, top-left). By using a custom MATLAB script to measure the Mal3- GFP binding area on the microtubules (Reid et al., 2017), we found that the fraction of microtubule area bound by Mal3-GFP was ∼2.7-fold lower in the presence of Eribulin as compared to the control experiments (Fig. 3D, bottom-left and Fig. S4A; p<<0.0001, Tukey’s Post-Hoc test after ANOVA). Similar results were observed for EB1-GFP (Fig. S4B, C; p<<0.0001, Tukey’s Post-Hoc test after ANOVA).

These results suggest that Eribulin binds to protofilament-edge sites along the damaged GTP-analogue microtubules, thus blocking protofilament-edge binding access for EB1. If this is true, we predicted that, in experiments without damaged microtubules, and thus with few protofilament-edge sites along the microtubule length, Eribulin would not have a significant impact on EB1 binding to the microtubule lattice. Thus, we performed similar EB1/Mal3 binding experiments, but used untreated, undamaged GMPCPP microtubules. We found that there was no significant difference in Mal3-GFP binding to undamaged microtubules in the presence or absence of Eribulin, consistent with the idea that Eribulin binds exclusively to protofilament- edge sites (Fig. 3D, right; p=0.36, Tukey’s Post-Hoc test after ANOVA; similar to EB1 results, Fig. S4B, C). Strikingly, Mal3-GFP binding was similar for undamaged microtubules in the absence of Eribulin, and damaged microtubules in the presence of Eribulin, suggesting that Eribulin efficiently blocked Mal3 binding to protofilament edge sites on the damaged microtubules (Fig. 3D, center-bottom, p=0.43, Tukey’s Post-Hoc test after ANOVA; EB1 results Fig . S4A).

### Eribulin suppresses Mal3-GFP tip tracking on dynamic microtubule plus-ends in cell-free experiments

Since Eribulin binds to protofilament-edge sites (Fig. 3B) (Doodhi et al., 2016; Jordan et al., 2005; Smith et al., 2010), and suppresses Mal3-GFP binding exclusively on damaged microtubules that have an increased density of protofilament-edge sites (Fig. 3D and Fig. S4A), we conclude that Eribulin blocks EB1/Mal3 binding at protofilament-edge sites (Fig. 3A).

Therefore, we reasoned that Eribulin could serve as an effective tool to test our simulation predictions regarding the role of protofilament-edge binding in EB1 tip tracking. Specifically, the simulation predicts that reducing the protofilament-edge on-rate by 4-fold will lead to a dramatic loss in Mal3-GFP intensity at the tips of dynamic microtubules (Fig. 4A, center), with a slightly larger loss in Mal3-GFP intensity when the protofilament-edge on-rate is reduced by 16- fold (Fig. 4A, right).

**Figure 4.**
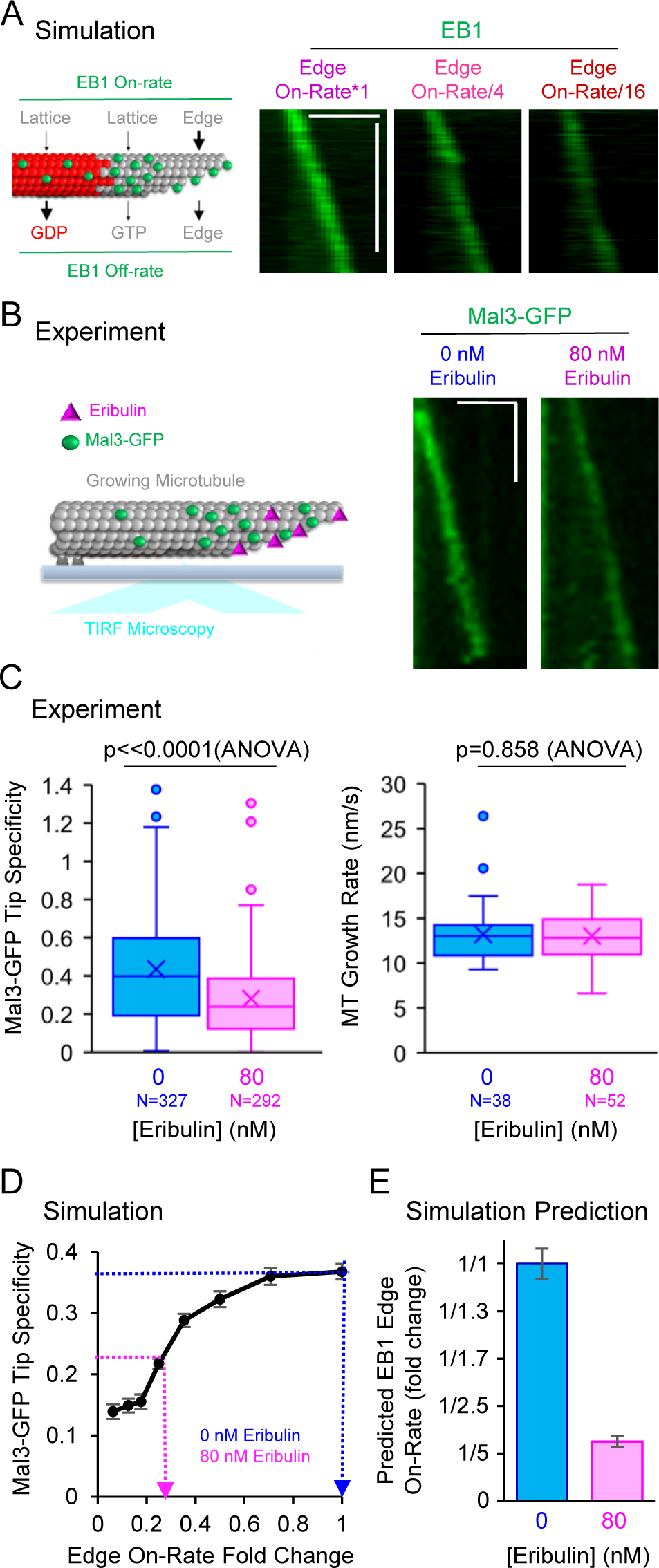
In cell-free experiments, Eribulin disrupts EB1 tip tracking by blocking EB1 access to protofilament-edge sites. (A) Left: Cartoon depicting relative differences in on-and off-rates in the simulation. Right: Simulated kymographs of EB1-GFP at growing microtubule tips, with decreasing protofilament-edge on-rates (Scale bars: horizontal: 2 µm, vertical: 60 s). (B) Left: Cartoon depicting experimental setup for dynamic microtubules with Mal3-GFP, in the presence of Eribulin, and visualized using TIRF microscopy. Right: Experimental kymographs of Mal3-GFP tip tracking along dynamic microtubules in the presence of 0 or 80 nM Eribulin (Scale bars: horizontal: 2 µm, vertical: 60 s) (C) Left: Eribulin reduces the Mal3-GFP Specificity in cell-free experiments (One Way ANOVA, p<<0.0001). Right: The presence of Eribulin did not significantly impact the microtubule growth rate in cell-free experiments (One-way ANOVA, p=0.858). (D) Simulation prediction for the median Mal3-GFP Specificity as a function of the fold-change in the protofilament-edge on-rate. For the protofilament-edge on-rate, a fold-change value of 1 is equivalent to the baseline simulation parameter value of 2.4x10^-3^ nM^-1^ sites^-1^ s^-1^. Each dotted line corresponds to the experimental median Mal3-GFP specificity for a given Eribulin concentration (Error bars: standard error of the median). E) Using the look-up table in panel D, a simulation prediction was generated for the fold-change in protofilament- edge on-rate in the presence of Eribulin. Error bars represent Q1 to Q3 of bootstrapped experimental median Mal3-GFP Specificity, as described in Methods.

To test this idea, we performed a cell-free assay in which dynamic microtubules were grown from stabilized seed templates in the presence of Mal3-GFP, and with or without Eribulin (Fig. 4B, left). Mal3-GFP at the tips of growing microtubules was visualized using TIRF microscopy (Fig. 4B, right). We observed that the presence of Eribulin led to a reduced intensity of Mal3- GFP at the growing microtubule plus-end (Fig. 4B, right), similar to simulation predictions for a reduced protofilament-edge Mal3-GFP on-rate (Fig. 4A, right).

To evaluate Mal3-GFP localization to growing microtubule plus-ends, we estimated the Mal3- GFP “specificity” in each case. Here, we defined Specificity (*S*) as:

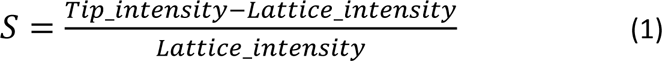

where the *Tip_intensity* was the summed Mal3-GFP intensity within a 4x4 pixel box at the growing microtubule tip, and the *Lattice_intensity* was the summed Mal3-GFP intensity in a 4x4 pixel box on the microtubule lattice. By definition, a Specificity value of *S*=0 means that the Mal3-GFP intensity at the growing microtubule tip is equal to the Mal3-GFP intensity on the lattice, away from the tip, and therefore Mal3-GFP is not localized at the growing microtubule plus-end.

Using the Specificity metric, we observed a ∼40% reduction in Mal3-GFP localization to the growing microtubule tip in 80 nM Eribulin as compared to control experiments in the absence of Eribulin (Fig. 4C, left, p<<0.0001, One-way ANOVA). Furthermore, the microtubule growth rate remained constant in the presence and absence of Eribulin, indicating that the loss in Mal3-GFP specificity was not due to a reduction in GTP-cap size (Fig. 4C, right, p=0.86, One-way ANOVA).

To predict the magnitude of protofilament-edge site blocking caused by Eribulin, we generated a look-up table from our simulations, in which we plotted the predicted Mal3-GFP specificity as a function of the fold-reduction in the protofilament-edge on rate (Fig. 4D, Fig. S4E). Using this look-up table, we estimated that an 80 nM Eribulin treatment led to a ∼4-fold reduction in the protofilament-edge on-rate (Fig. 4E).

### In cells, Eribulin reduces EB1 localization at growing microtubule ends

Finally, we asked whether protofilament-edge site blocking via Eribulin could suppress EB1 tip tracking of growing microtubules in cells. Thus, increasing concentrations of Eribulin were added to fresh imaging buffer in dishes of cultured LLC-PK1 cells that stably expressed EB1-GFP (Piehl & Cassimeris, 2003), and allowed to incubate for one hour. We note that a range of low Eribulin concentrations were selected (0-50 nM), in order to maintain a constant microtubule growth rate (See Fig. S5F, Movie S9 for higher Eribulin concentration that disrupts microtubule growth rate). The cells were imaged using confocal microscopy (Fig. 5A, Movies S7, S8), and the EB1-GFP Specificity (*S*) was measured in each case (Eqn. 1; See Methods).

**Figure 5.**
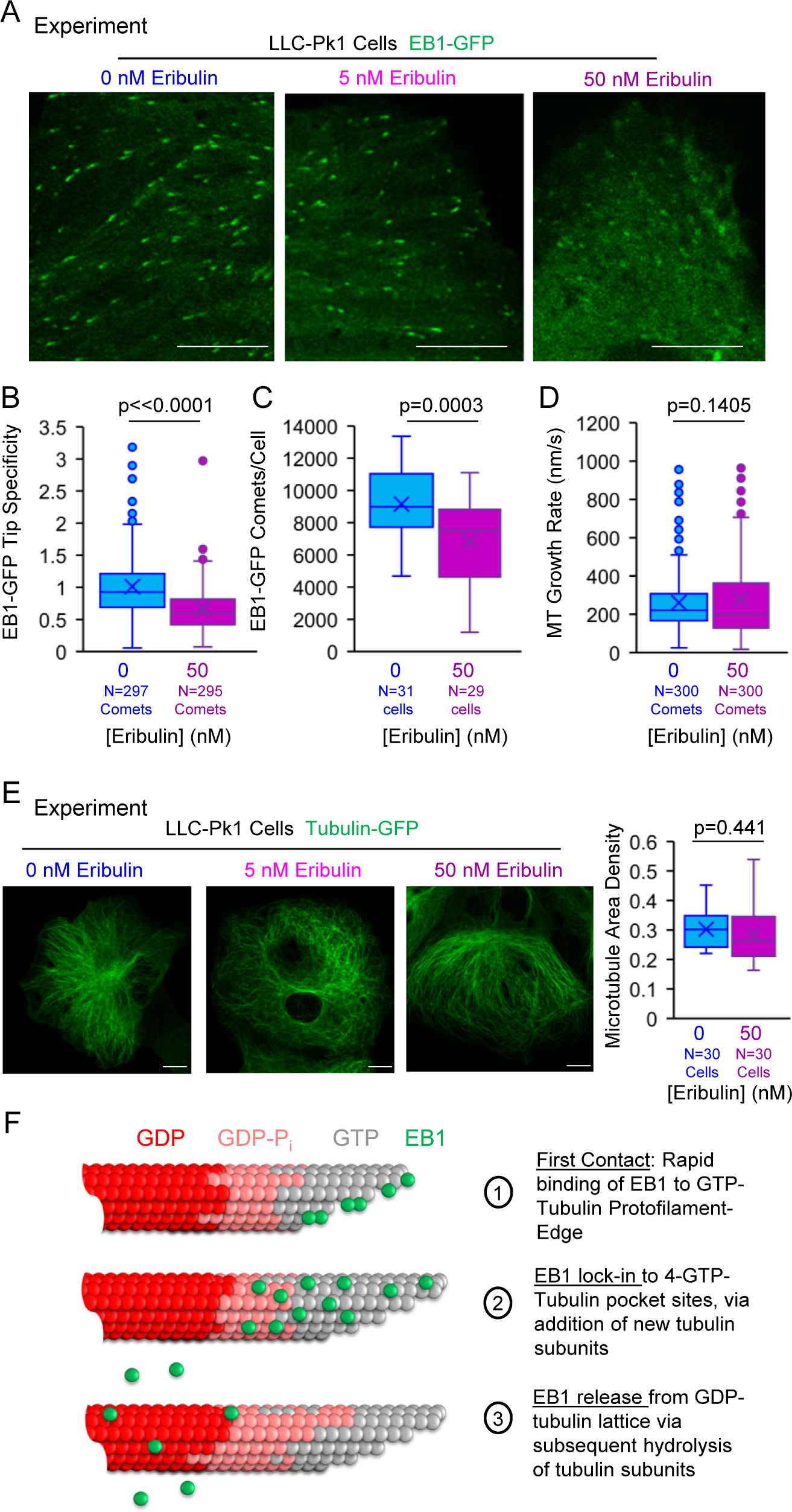
Eribulin disrupts EB1 tip tracking in cells. (A) EB1-GFP comets in live LLC-PK1 cells treated with either DMSO control (left), 5 nM Eribulin (middle), or 50 nM Eribulin (right) (Scale bar: 10 µm). (B) EB1-GFP Tip Specificity for individual comets in cells treated with 0 or 50 nM Eribulin (p<<0.0001, t-test). (C) Average number of EB1-GFP comets per cell in cells treated with 0 or 50 nM Eribulin (See Methods, (Applegate et al., 2011); p=0.0003, t-test). (D) Growth rate of individual microtubules in cells treated with either 0 or 50 nM Eribulin (p=0.141, Mann-Whitney U test). (E) Left: Tubulin-GFP in cells treated with either DMSO control, 5 nM Eribulin, or 50 nM Eribulin (Scale bar 10 µm). Right: Quantification of microtubule density in cells treated with 0 or 50 nM Eribulin (p=0.553, t-test; See Methods). (F) Model for EB1 tip tracking: 1) EB1 rapidly binds to protofilament-edge sites, with a slower on-rate to 4-tubulin closed lattice binding sites. 2) Incorporation of new tubulin subunits “locks in” EB1 bound at protofilament-edge sites as the binding pocket transitions from a protofilament-edge site to a 4-tubulin closed-lattice site. 3) As the GTP-tubulin hydrolyzes to GDP-tubulin, the EB1 dissociates from the GDP closed- lattice.

In the presence of 50 nM Eribulin, the EB1 specificity was reduced by ∼34% relative to the DMSO control (Fig. 5B, p<<0.0001, t-test; 5 nM data Fig. S5A). Of note, if the EB1-GFP Specificity drops down to zero, the EB1-GFP intensity at the growing microtubule tip would match the EB1-GFP intensity along the length of the microtubule, and so an EB1-GFP tip comet would not be apparent at all, leading to a reduction in the number of comets observed per cell. Consistent with this idea, we observed a ∼25% reduction in the average number of EB1-GFP comets per cell in 50 nM Eribulin relative to the DMSO control (Fig. 5C, p=0.0003, t-test; 5 nM Eribulin data, Fig. S5B). Similar to our cell-free assays, this disruption in EB1-GFP tip tracking occurred despite little or no change in the cellular microtubule growth rate (Fig. 5D, Fig. S5D; 5 nM Eribulin data,

Fig. S5C; see Methods (Applegate et al., 2011)) or overall microtubule density in the cell (Fig. 5E; 5 nM Eribulin data, Fig. S5E).

## Discussion

In this work, we developed a molecular-scale computational simulation that incorporated both tubulin and EB1 on-off dynamics. Our simulation predicted that binding of EB1 to protofilament-edge sites contributes to efficient tip-tracking of EB1 at growing microtubule plus-ends. To test this prediction, we used Eribulin, a chemotherapy drug, which binds to protofilament-edge sites on microtubules. We found that Eribulin blocks EB1 binding at protofilament-edge binding sites, and importantly, this blocking of EB1 binding to protofilament-edge sites led to a dramatic reduction in EB1 tip tracking in cells, and in cell-free assays. Thus, we conclude that the rapid binding of EB1 to protofilament-edge sites facilitates the tip tracking of EB1 at growing microtubule ends.

Based on our results, we have developed a working model for tip tracking in which EB1 binds at a rapid rate to GTP-tubulin protofilament-edge sites at the tip of the microtubule (Fig. 5F, 1).

We note that the off-rate of EB1 from these partial binding sites is likely higher than the off-rate of EB1 from closed 4-tubulin pocket binding sites. However, as the microtubule grows and new GTP tubulin dimers are incorporated, EB1 bound at protofilament-edge sites become “locked in” to their 4-tubulin pocket binding sites by the addition of new tubulin dimers, leading to low EB1 off-rates (Fig. 5F, 2). Thus, EB1 accumulates on the GTP-cap at the growing microtubule end, with a peak EB1 position slightly distal from the growing microtubule tip, as has been previously reported (Maurer et al., 2014; Roth et al., 2019). Finally, upon the hydrolysis of GTP-tubulin to GDP-tubulin, the affinity of EB1 for the GDP-tubulin subunits is reduced, and EB1 dissociates from the microtubule (Fig. 5F, 3). Therefore, the higher affinity of EB1 for GTP relative to GDP tubulin also contributes to EB1 localization at growing microtubule ends. This model explains how Eribulin, which binds primarily to protofilament-edge sites, could disrupt EB1 tip tracking.

Recent work has demonstrated that growing microtubule tips are less homogeneous than previously thought, such that growing microtubule tips exhibit a wide range of protofilament lengths between the leading and lagging protofilaments, both in cells and in cell-free experiments (Atherton et al., 2018; Cleary & Hancock, 2021; C. E. Coombes et al., 2013; Gudimchuk et al., 2020; Guesdon et al., 2016; Igaev & Grubmüller, 2022). Thus, protofilament- edge sites that are present due in part to a wide range of protofilament lengths at growing microtubule tips could contribute to the localization of protofilament-edge binding proteins, such as EB1, to growing microtubule ends.

Recently published work provides support for the importance of EB1 protofilament-edge site binding in the efficiency of EB1 tip tracking. Specifically, by using the microtubule polymerase protein XMAP215 in cell-free experiments, Farmer et al increased the range of protofilament lengths between the leading and lagging protofilaments (increased “tip tapering”) (Farmer et al., 2021). Importantly, Farmer et al also observed a dramatic increase in EB1-GFP intensity at the growing microtubule tip with increasing XMAP215-induced tip taper at the microtubule plus-ends, independent of microtubule growth rate (Farmer et al., 2021). We note that increased tip taper would correspond to an increase in the average number of protofilament- edge sites at the growing microtubule end. Thus, XMAP215 could increase the efficiency of EB1 tip tracking by adding new protofilament-edge sites to the growing microtubule plus-end.

Our stochastic simulation robustly recapitulated experimental EB1 tip tracking, including previously reported localization of EB1 relative to the growing microtubule plus-end, and dependencies of EB1 tip tracking on the GTP-tubulin hydrolysis rate and microtubule growth rate. The model was robust over a wide range of model parameter values, as even the most sensitive parameters had at least an 8-fold range of acceptable values in which EB1 was clearly localized exclusively at the growing microtubule end (Fig. S1, S2). Future experiments could use newly described methods to examine the relationship between the EB1 on and off rates relative to tubulin on and off rates, to more closely examine our new model for rapid EB1 dynamics at protofilament-edge sites, combined with a slower, hydrolysis-dependent EB1 off-rate from 4-tubulin pocket sites along the GTP-cap (Mickolajczyk et al., 2019). Further, microtubule targeting drugs that suppress the kinetics of tubulin assembly at the growing microtubule plus-end, such as Taxol, could alter EB1 tip tracking by slowing the capture and “lock in” of EB1 to 4-tubulin pocket binding sites by the addition of new tubulin dimers (Fig. 5F, 2), an idea that could be explored in future work.

Consistent with our observations, multiple groups have observed a disruption in EB1 tip tracking with Eribulin treatment in cells (Chanez et al., 2015; Doodhi et al., 2016; O’Rourke et al., 2014). Furthermore, western blotting has been used to demonstrate that this Eribulin- dependent suppression of EB1 tip tracking in cells was not due to a reduction in EB1 expression (Chanez et al., 2015). Previous work using a cell-free system has also described a growth-rate independent effect of Eribulin on the disruption of EB3 tip tracking, similar to our observations (Doodhi et al., 2016). In this work, the authors speculate that the origin of this tip-tracking suppression was the tapering of the microtubule tip, leading to “split” EB3 comets and thus fewer EB3 canonical binding sites (Doodhi et al., 2016). However, we rarely observed split comets in our experiments, likely due to our slower experimental microtubule growth rate (∼15 nm/s in our assay vs ∼50 nm/s in (Doodhi et al., 2016)).

In conclusion, we find that protofilament-edge sites are an important contributing factor for proper EB1 tip tracking along growing microtubule ends, and that EB1 tip tracking is suppressed by blocking the protofilament-edge sites at growing microtubule ends. Therefore, using drugs or proteins to alter the number of exposed protofilament-edge sites at the growing microtubule tip may provide a new mechanism to regulate EB1 tip tracking in cells.

## Materials and Methods

### Cell lines

The LLC-Pk1 cell line expressing EB1-GFP was a gift from Dr. Patricia Wadsworth, and the cell line expressing GFP-Tubulin was a gift from Dr. Lynne Cassimeris. The identity of the cell lines (non-human) were authenticated by microscopy observation and analysis.

### Culture, Imaging, and drug treatment of LLC-PK1 cells

The LLC-Pk1 cell lines were grown in Optimem media (ThermoFisher #31985070), 10% fetal bovine sera + penicillin/streptomycin at 37°C and 5% CO_2_. Cells were grown in 14-mm glass bottom dishes for visualization by microscopy. For drug treatments, the Optimem media was replaced with CO_2_-independent imaging media. Drug dissolved in DMSO, or only DMSO as a control, was added to the final concentration. Cells were incubated with the drug/DMSO for 1 hour prior to imaging and were imaged at 37 °C. Cells were imaged with a laser scanning confocal microscope (Nikon Ti2, 488nm laser line) fitted with a 100x oil objective (Nikon N2 Apochromat TIRF 100X Oil, 1.49 NA), which allowed for a 0.16 μm pixel size.

### LLC-PK1 comet image analysis

To analyze EB1-GFP comets in LLC-Pk1 cells, the brightest comets in each few frames were cropped, and their direction was noted. These comets were then not cropped in later frames to ensure each comet was only analyzed once during its lifetime. To determine the EB1-GFP Specificity (Eqn. 1), a custom MATLAB script was written that allowed the user to pick the brightest point of the comet and then a point on the microtubule lattice, behind the comet.

Then, a 4 X 4 pixel box was summed at the brightest point of the comet (*Tip_intensity*) and a separate 4 X 4 pixel box was summed at the point behind the comet (*Lattice_intensity*). Then, Equation 1 was used to calculate the EB1 Specificity (*S*). This method was repeated for many comets across multiple cells from three biological replicates for each condition. The EB1 specificity was compared across treatment conditions using a Student’s T-Test.

### LLC-Pk1 Microtubule growth rate analysis

To analyze EB1-GFP comet velocity, which was used as a proxy for the microtubule growth rate, we employed analysis software from the Danuser lab (Applegate et al., 2011). In short, we collected multiple videos of LLC-Pk1 cells treated with DMSO or Eribulin from three separate biological replicates and loaded them into the Danuser code software, using constant parameters for thresholding and water shedding. We then allowed the program to identify, link, and track comets over time, which provided us with EB1 comet velocities across multiple cells. We next cut off any outlier values greater than 1 um/s, which were likely artifacts from the analysis software. Then, to match sample sizes in the Specificity measurements, 300 growth rate values were randomly selected from the thousands of comets the software detected, and were used to compare microtubule growth rates across conditions. The software also provided information about the number of comets identified in each cell, which was also used in our analysis. The growth rates were statistically analyzed using a Mann-Whitney U test while the number of comets were statistically analyzed using a Student’s T-Test.

### LLC-PK1 Microtubule density analysis

To determine the microtubule density in LLC-Pk1 cells treated with Eribulin or a DMSO control, LLC-Pk1 cells overexpressing Tubulin-GFP were subjected to Eribulin or DMSO treatment, followed by imaging with a confocal microscope. Z-stacks were acquired across the volume of these cells. Then, maximum Z-projections were created, followed by analysis of the area of the microtubules divided by the area of the cell, which was performed with a custom MATLAB script (Goldblum et al., 2021). Finally, this normalized value was obtained for multiple cells across each condition from three separate biological replicates and compared using a One-Way ANOVA.

### Simulation Methods

The MATLAB scripts that were used in the manuscript are available on the Gardner lab website. https://cbs.umn.edu/gardner-lab/software-links.

The computational simulation combined both tubulin molecule assembly, and EB1 binding and unbinding.

The microtubule assembly portion of the simulation is based in part on work from the Goodson lab (Margolin et al., 2012), as well as prior work from the Gardner lab (C. E. Coombes et al., 2013). Briefly, this model allows microtubule protofilaments to grow independently via the addition of individual tubulin subunits, and to form and break lateral bonds with neighboring protofilaments once individual tubulin subunits are longitudinally bound to the microtubule.

The lateral bonding and breaking parameters were kept consistent throughout all simulations. See Table S1 for simulation parameters.

**Table S1:** Simulation Parameters: Microtubule Assembly

| Parameter | Description | Value | Reference |
| --- | --- | --- | --- |
| [GTP-tub] | Free GTP-Tubulin concentration | $3 \times 10^{-6} - 15 \times 10^{-6} \text{ M}$ | Matched to different experiments to have similar growth rates as experiment |
| $k_{\text{on,PF}}$ | Tubulin on-rate constant | $1.25 \mu\text{M}^{-1} \text{ s}^{-1} \text{ pf}^{-1}$ | (Margolin et al., 2012) |
| $k_{\text{off, GTP}}$ | Tubulin off-rate when GTP dimer below | $0.02 \text{ s}^{-1}$ | (Margolin et al., 2012) |
| $k_{\text{off, GDP}}$ | Tubulin off-rate when GDP dimer below | $20 \text{ s}^{-1}$ | (Margolin et al., 2012) |
| $k_{\text{hyd, GTP}}$ | Hydrolysis rate constant | $0.725 \text{ s}^{-1}$ | (Margolin et al., 2012; Reid et al., 2016) |
| $k_{\text{lateral Bond Formation}}$ | Formation rate for a lateral bond between protofilaments | $100 \text{ s}^{-1}$ | (Margolin et al., 2012) |
| $k_{\text{lateral bond break TT}}$ | Breakage Rate for lateral bond between two GTP dimers | $70 \text{ s}^{-1}$ | (Margolin et al., 2012) |
| $k_{\text{lateral bond break TD}}$ | Breakage rate for lateral bond between a GTP and GDP dimer | $90 \text{ s}^{-1}$ | (Margolin et al., 2012) |
| $k_{\text{lateral bond break DD}}$ | Breakage rate for lateral bond between two GDP dimers | $600 \text{ s}^{-1}$ | (Margolin et al., 2012) |
| $k_{\text{lateral bond break TT Seam}}$ | Breakage Rate for lateral bond two GTP dimers at the seam | $140 \text{ s}^{-1}$ | (Margolin et al., 2012) |
| $k_{\text{lateral bond break TD Seam}}$ | Breakage Rate for lateral bond between a GTP and GDP dimer at the seam | $180 \text{ s}^{-1}$ | (Margolin et al., 2012) |
| $k_{\text{lateral bond break DD Seam}}$ | Breakage Rate for lateral bond between two GDP dimers at the seam | $1200 \text{ s}^{-1}$ | (Margolin et al., 2012) |
| $\pi_{\text{break}}$ | Correction factor for the lateral bond breakage rate if the neighboring PFs all have lateral bonds | 5.4 | (Margolin et al., 2012) |

In the EB1 binding and unbinding portion of the simulation, the model allows for EB1 to bind and dissociate from the microtubule independently of tubulin addition/dissociation. Here, EB1 could bind to protofilament-edge sites and closed-lattice sites with differing on-rates, and then dissociated from GTP and GDP binding sites with differing off-rates. See Table S2 for simulation parameters.

**Table S2:**
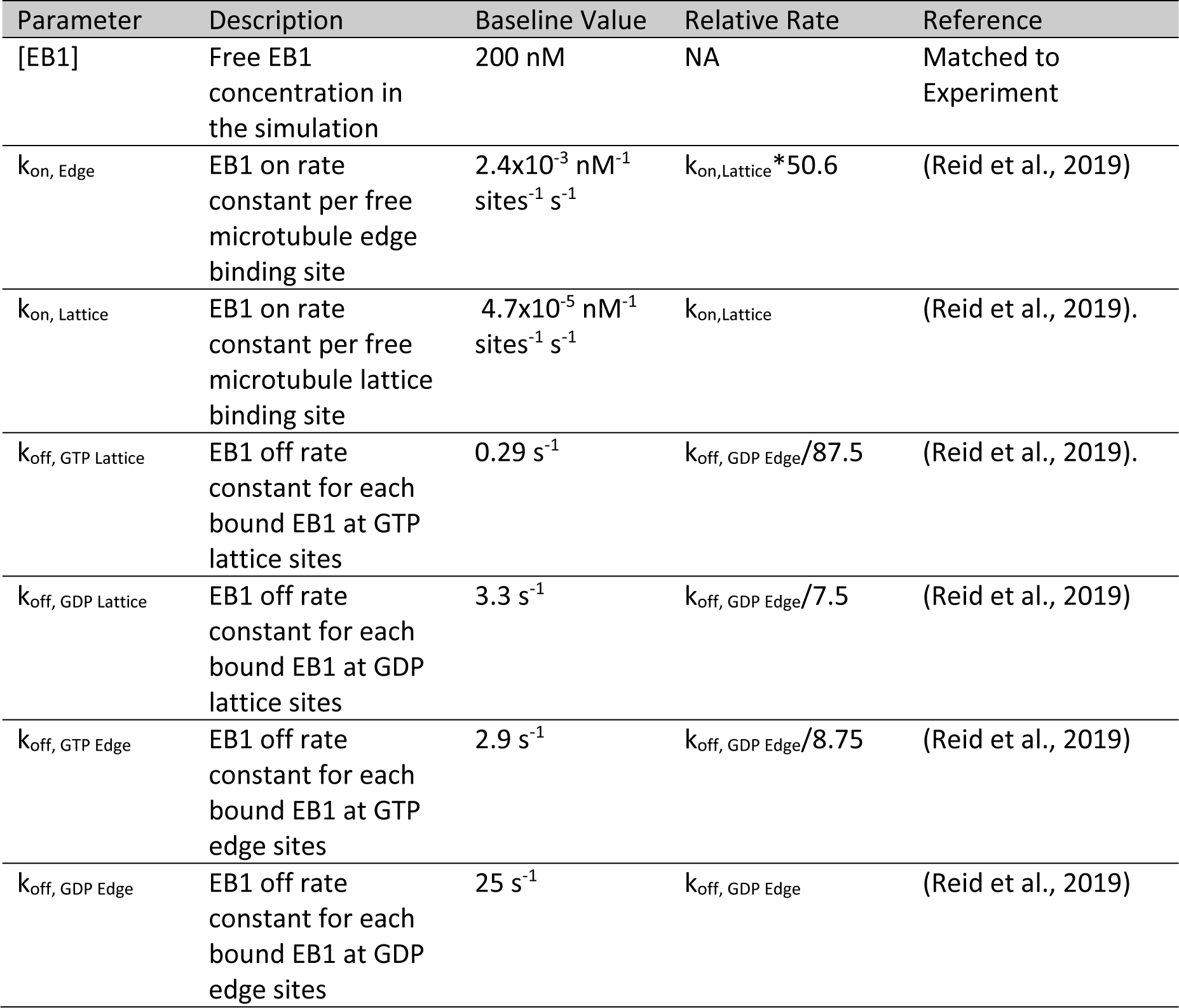
Simulation Parameters: EB1 Dynamics

At the start of every time step in the simulation, total execution time was calculated for each potential event, which included EB1 association/dissociation, tubulin association/dissociation, and lateral bond formation/breakage between protofilaments, using the equation:

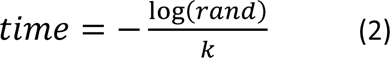

where *time* is the total execution time required for each potential event, *rand* represents the built-in MATLAB function that generates a uniformly distributed random number between 0 and 1, and *k* is the single-molecule rate constant for each potential event.

Then, once all the times were calculated for each potential event, the simulation executed only the one event with the shortest time.

After every time step in the simulation, every tubulin dimer’s state was recorded (GTP versus GDP), and every single potential EB1 binding site state was also recorded (GTP vs GDP; Edge vs Lattice; EB1 bound or not bound). With each step of the simulation, these values were updated based on what happened during that step (eg, tubulin association/dissociation, tubulin hydrolysis, EB1 binding/dissociation, lateral bond breakage/formation). Every thousand steps of the simulation (which averaged between 0.5 and 2 seconds of real time in the simulation), the length of each microtubule protofilament and the position of every bound EB1 were stored for later use. This was repeated until the simulation ended, which in general tended to be somewhere between 6x10^4^and 2x10^5^ steps, depending on which simulation experiment was being performed.

### Creating model-convolved images from the simulation

To visualize the output from the simulation, 15% of the tubulin dimers were randomly labelled with a red fluorophore and plotted on an image. Then, all EB1 positions were labelled with a green fluorophore and plotted. Next, random, simulated noise was added to the image. Finally, these points were convolved using a point spread function as would be observed using a 100X objective on our TIRF microscope (Demchouk et al., 2011; Gardner et al., 2010; Reid et al., 2019) This model-convolution process was performed using the stored data from every 1000 steps in each simulation, and allowed for the generation of simulated movies in which the simulated EB1 localization along growing microtubules could be observed.

### Analyzing line scans of simulated Images for Peak EB1 position and tip/lattice intensity

To quantify EB1 tip tracking in the simulation, line scans along the length of simulated microtubules were obtained using ImageJ. Then, each image was aligned along the peak EB1 position. Next, at least five separate simulation runs were averaged. To determine the end of the microtubule, the position where the intensity of the microtubule channel (red) was halfway between its maximum value and its minimum value was deemed as the microtubule end, and denoted with a relative position of zero nanometers. From here, the peak EB1 position could be determined by finding its relative position with respect to the microtubule end at position 0. Of note, in all simulated data that used line scans, N refers to the number of individual simulation runs, except for the peak position values. In this case, N refers to the average value from 5 simulation runs, so the N=5 means that the simulation was performed 25 times and averaged in sets of 5 before being used. To determine the tip/lattice intensity values, the maximum EB1 intensity/pixel at the tip was divided by the median EB1 intensity/pixel along the lattice, which 768-1024 nanometers distal of the microtubule plus end.

### Analyzing line scans of simulated images for microtubule tapering

To quantify the microtubule tip standard deviation in the simulation (Fig. S1A), the simulated images used for the EB1 peak position analysis were loaded into a previously described program to measure tip tapering (Demchouk et al., 2011). In short, the user defined points along the start and end of the microtubule lattice and then the intensity of the microtubule across this distance was measured. Then, this intensity was fit to a Gaussian survival curve to determine the tip standard deviation of the microtubule (which includes the microscope point spread function as well as the underlying protofilament length standard deviation). Of note, each data point is the standard deviation for a single simulation run. Finally, the average of this value was compared to a previously reported average tip standard deviation value (Maurer et al., 2014). Of note, outliers were removed if their fitted tip standard deviation was greater than two times the true microtubule taper obtained from the simulation protofilament lengths.

### Simulation parameter testing

To determine the robustness of tip tracking in our simulation, we ran a series of simulations in which, in each case, all parameters but one were held constant and then this one parameter was altered in 2-fold increments. The range for this parameter testing spanned from 1/16 to 16 fold times the baseline parameter value for each key parameter in the simulation. Ten simulation runs were performed for each parameter set. In each case, a simulated movie was generated, and a line scan was obtained, as described above. To determine the best fit for a given parameter across the range we tested, we calculated the sum of absolute errors between the simulated EB1 line scan profile and the line scan profile for Mal3-GFP from data in the literature (Bieling et al., 2007), and selected the parameter value with the smallest sum of absolute errors as the best-fit parameter value.

### Simulation turning off edge binding and turning back on

To remove protofilament-edge binding partway through the simulation, we added a condition inside our script that set the protofilament-edge on-rate to zero after 25,000 steps in the simulation had been performed. Then, we added another condition which changed the edge on rate back to what it was at the start of the simulation after another 35,000 steps of the simulation had occurred. A similar simulation was run for the closed-lattice binding case.

### Labelling of 20-nm gold beads with Eribulin

20nm gold particles were coated with Eribulin by using an NHS-activated gold nanoparticle conjugation kit (Cytodiagnostics CGN5K-20-1), following the kit instructions, except that a greater molar excess of Eribulin was used than was suggested by the instructions for antibody labeling, as follows: 4.3 μL of 10 mM Eribulin in DMSO was added to 47 μL of the kit’s dilution buffer. 60 μL of the kit reaction buffer was then added. 90 μL of this solution was then added to the vial of lyophilized gold particles and incubated at room temperature for 2 hrs, triturating every 15 min. 10 μL of the quencher reagent was added then centrifuged at 6000xg for 30min. The gold particle pellet was washed with 400ul of storage buffer (20mM Tris pH8 / 150mM NaCl / 1% BSA / 0.025% tween-20), centrifuged at 8000 xg for 10min., and then particles were resuspended in 400 μL of storage buffer, aliquoted, frozen with liquid nitrogen and stored at - 80°.

### Tubulin purification and labelling

Tubulin was purified and labeled as previously described (Gell et al., 2010).

### Preparation of GTP-analogue microtubules

GMPCPP microtubules were prepared as previously described (Gell et al., 2010). In short, 3.9 µM rhodamine labelled tubulin was incubated with 1 mM GMPCPP with 1.1 mM MgCl_2_ in BRB80 for 5 minutes on ice. Then, the solution was incubated at 37 °C for 2 hours or overnight. The GMPCPP microtubules were then spun with an airfuge (Beckman Coulter, 20psi, 5min) and further stabilized in 10 μM Taxol in BRB80. These microtubules were stored at 37 °C and used in experiments within 4 days after preparation. These microtubules were used as seeds for dynamic microtubule experiments and were used for EB1/Mal3 binding along undamaged GMPCPP microtubules.

### Negative stain TEM for Eribulin binding to microtubules

To perform Negative Stain TEM of microtubules, GDP microtubules were made by incubating 66 µM of free tubulin with 1.2 mM GTP and 4.8 mM MgCl_2_ first on ice for 5 minutes, followed by 30 minutes at 37 °C. Next, this solution was diluted 100-fold in a 10 μM taxol in BRB80 solution and allowed to sit at room temperature overnight.

The GDP microtubules were then incubated with Eribulin bound to 20 nM gold beads at room temperature for one hour. Next, 300-mesh carbon coated copper grids were warmed for 5-10 minutes on a hotplate. After warming, 10 µL of the microtubule and Eribulin-bound gold bead solution was added to a grid and allowed to sit for one minute, followed by adding a few small drops of 0.5% uranyl acetate. The grid was immediately tilted to allow any excess uranyl acetate to drop off, and then excess uranyl acetate was wicked away with a Kim wipe. The grid was then allowed to sit for 5-10 minutes before placing back in storage until imaging. The samples were imaged using an FEI Tecnai T12 Transmission electron microscope.

### Analysis of TEM for Eribulin-bound beads on microtubules

To quantify the binding of Eribulin-bound gold beads to different microtubule structures (closed-lattice or protofilament-edges), 1.75 μm X 1.75 μm fields of view were captured using the FEI Tecnai T12 Transmission electron microscope. In each field of view, the number of beads bound to the microtubule lattice was divided by the combined length of all microtubules in the field of view. Likewise, the number of beads bound to microtubule protofilament-edges was divided by the combined length of all protofilament-edge binding sites at microtubule ends in the field of view. To be consider bound, the bead needed to partially overlap with the microtubule.

### Purification of EB1-GFP

Plasmid pETEMM1-HIS6x-tev-EB1-GFP was transformed into Rosetta (DE3) pLysS E. coli, grown in 10 ml of LB+cam+kan at 37° overnight, and then subcultured into 1L of the same media, mixing at 37° for 2 hr to an A600 of 0.44. IPTG was then added to 2 mM and the culture was mixed at 18° for 14 hr. The culture was centrifuged 30 min. at 4°, 4400 xg. The cell pellet was resuspended into 12 ml of PBS/0.1% tween-20/5 mM B-mercaptoethanol/1 mg/ml lysozyme and protease inhibitors (1 mM AEBSF/10 mM pepstatin A/0.3 mM aprotinin/10 uM E-64), mixed at 4°, 2 hr., and then sonicated on ice at 80% power, 50% duty, for 10 cycles of 1 min. on, 1 min. off. The lysate was centrifuged at 4°, 1 hr., 18000 xg. The supernatant was passed through 2 ml of Talon Metal Affinity Resin and the resin was sequentially washed with five column volumes Buffer A (50 mM sodium phosphate pH7.5/300 mM KCl/10% glycerol/5 mM B-mercaptoethanol). Next, the column was washed with 95% buffer A/5% bufferB (Buffer A/300 mM imidazole), followed by a wash with 90% A/10% B, and finally 85% Buffer A/15% Buffer B. All buffers had the protease inhibitors 1 mM AEBSF, 10 uM pepstatin A and 10 uM E-64. Protein was eluted with 1 ml fractions of 100% Buffer B + above protease inhibitors. Elution fractions were analyzed on coomassie and western blot. Relevant fractions were combined and dialyzed against Buffer A overnight at 4°, then 2 hr. at 18° in the presence of HIS6-taggedTEV protease (Expedeon TEV0010) at a 1:100 w/w protein ratio). Dialysate was then mixed with additional Talon resin to remove cleaved HIS6x tag.

### Purification of Mal3-GFP

The pETMM11-HIS6x-Mal3-GFP plasmid with a TEV cut site after the His6x tag was a kind gift from Dr. Thomas Surrey. The plasmid was transformed into Rosetta (DE3) pLysS E. coli and grown in 800mL of LB+kan+cam at 37°C to an OD of approximately 0.4. To induce protein expression, IPTG was added to 0.2mM and the culture was mixed at 14°C for 16hr. Cells were centrifuged (30min., 4°C, 4400xg) and resuspended in 25mL lysis buffer (50mM Tris pH7.5, 200mM NaCl, 5% glycerol, 20mM imidazole, 5mM β-mercaptoethanol, 0.2% triton X-100), protease inhibitors (1mM PMSF, 10μM Pepstatin A, 10μM E-64, 0.3μM aprotinin), and DNAse I (1U/mL). The cell suspension was sonicated on ice (90% power, 50% duty, 6x1min). Cell lysates were centrifuged (1h, 4°C, 14000 xg) and the soluble fraction was passed through 1mL of Talon Metal Affinity Resin (Clontech #635509). The resin was washed for four times with 4mL Wash Buffer (50mM Tris pH7.5, 500mM NaCl, 5% glycerol, 20mM imidazole, 5mM β- mercaptoethanol, 0.1mM PMSF, 1μM Pepstatin A, 1μM E-64, 30nM aprotinin) for 5min each.

Protein was eluted from the resin by mixing with 1mL of Elution Buffer (50mM Tris pH 7.5, 200mM NaCl, 250mM imidazole, 0.1mM PMSF, 1μM Pepstatin A, 1μM E-64, 30nM aprotinin) for 15min followed by slow centrifugation through a fritted column to retrieve eluate. To cleave the HIS6x tag, 10 units of GST-tagged TEV enzyme (TurboTEV, #T0102M, Accelagen) and 14mM β-mercaptoethanol were added and the eluate was dialysed into Brb80 overnight at 4°C. To remove the TEV enzyme, the dialysate was mixed with 100ul of glutathione-sepharose (GE Healthcare #17-0756-01) for 30 min. at 4°C and spun (1min, 2000xg). The Mal3-GFP protein was quantified by band intensity on a coomassie-stained SDS PAGE protein gel.

### Creation of TIRF microscopy Flow Chambers for cell-free assays

Imaging flow chambers were constructed as in Section VII of Gell et al. (2010), with the following modifications: two narrow strips of parafilm replaced double-sided scotch tape as chamber dividers: following placement of the smaller coverslip onto the parafilm strips, the chamber was heated to melt the parafilm and create a seal between the coverslips; typically, only three strips of parafilm were used, resulting in two chambers per holder. Chambers were prepared with anti-rhodamine antibody (Invitrogen A6397, RRID:AB_2536196) followed by blocking with Pluronic F127, as described in Section VIII of Gell et al. (2010), for dynamic microtubule experiments and for EB1/Mal3 binding to stabilized, CaCl_2_ damaged microtubules. To reduce background for the undamaged microtubule experiments, we used microtubules labeled with biotin and rhodamine and adhered them to a plasma treated coverslip with an anti-rhodamine and streptavidin mix. The coverslips were then blocked with 1 mg/mL PLL-PEG followed by 1 mg/mL casein. Microtubules were adhered to the chamber coverslip, and the chamber was flushed gently with warm BRB80. The flow chamber was heated to 28°C using an objective heater on the microtubule stage, and then 3–4 channel volumes of imaging buffer were flushed through the chamber. Microtubules were imaged on a Nikon TiE microscope using 488 nm and 561 nm lasers sent through a Ti-TIRF-PAU for Total Internal Reflectance Flourescence (TIRF) illumination. An Andor iXon3 EM-CCD camera fitted with or without a 2.5X projection lens depending on the experiment was used to capture images with high signal to noise and small pixel size (64 nm or 160 nm respectively). Images were collected using TIRF with a Nikon CFI Apochromat 100 X 1.49 NA oil objective.

### Cell-free microtubule assay setup

For the damaged GTP-analogue microtubule assays, GMPCPP microtubules were introduced into a flow chamber as described above and allowed to incubate for 30 seconds to 3 minutes before flushing out any non-adhered microtubules with BRB80. Next, 40 µM CaCl_2_ in warmed BRB80 was introduced into the chamber and incubated for 3-5 minutes, until obvious degradation occurred to the microtubules. The chamber was then washed with multiple chamber volumes of warmed BRB80. Next, the chamber was washed with one chamber volume of prewarmed imaging buffer (20μg/mL glucose oxidase, 10μg/mL catalase, 20mM D-Glucose, 10mM DTT, 80μg/mL casein, 110 mM KCl and 1% tween-20). Finally, an EB1/Mal3 reaction mixture with or without 160 nM Eribulin (imaging buffer plus 123 nM Mal3-GFP or 165 nM EB1- GFP) was introduced to the chamber and allowed to incubate for 15 minutes. Images of hundreds of non-overlapping fields of view were collected and used for downstream analysis.

For undamaged GTP-analogue microtubule assays, GMPCPP microtubules were introduced into a flow chamber as described above, and allowed to incubate for 30 seconds to 10 minutes before flushing out any non-adhered microtubules with BRB80. Next, one chamber volume of prewarmed imaging buffer (20μg/mL glucose oxidase, 10μg/mL catalase, 20mM D-Glucose, 10mM DTT, 80μg/mL casein, 110 mM KCl and 1% tween-20) was added. Finally, a reaction mixture with or without 160 nM Eribulin (imaging buffer plus 123 nM Mal3-GFP or 165 nM EB1- GFP) was introduced to the chamber and allowed to incubate for 15 minutes. Finally, images of hundreds of non-overlapping fields of view were obtained and used for downstream analysis.

For the dynamic microtubule assays, GMPCPP microtubule seeds were introduced into a flow chamber as described above and allowed to incubate for 30 seconds to 3 minutes before flushing out any non-adhered microtubules with BRB80. Next, one chamber volume of prewarmed imaging buffer (20μg/mL glucose oxidase, 10μg/mL catalase, 20mM D-Glucose, 10mM DTT, 80μg/mL casein, 110 mM KCl and 1% tween-20) was added. Finally, a reaction mixture consisting of Imaging buffer plus 212 nM Mal3-mCherry, 11 uM of 12% green-labelled tubulin, and 1 mM GTP, with or without varying concentrations of Eribulin (0, 80, or 160 nM) was added. Time-lapse images were collected of dynamic microtubules growing from the GMPCPP stabilized seeds and were then used for quantification.

### Analyzing microtubule area bound by EB1-GFP/Mal3-GFP

To compare the binding of EB1-GFP or Mal3-GFP on undamaged and damaged microtubules in the presence and absence of Eribulin, the total length of green (EB1-GFP or Mal3-GFP) occupancy was divided by the total length of the red microtubules on each image (defined as EB1-GFP or Mal3-GFP ‘microtubule coverage’). This was accomplished by using previously described semi-automated MATLAB analysis code (Reid et al., 2017). Briefly, first, automatic processing of the red microtubule channel was used to determine the microtubule-positive regions, which then allowed for conversion of the red channel into a binary image with white microtubules and a black background. The green EB1-GFP/Mal3-GFP channel was then also pre- processed to smooth high-frequency noise and to correct for TIRF illumination inhomogeneity. The green channel threshold was then manually adjusted to ensure visualization of all EB1-GFP binding areas on each microtubule. Measurements of the total EB1-GFP or Mal3-GFP coverage area were then automatically collected from the identified microtubule regions. Finally, the total coverage area of EB1-GFP or Mal3-GFP was divided by the total microtubule area in each field of view. This experiment was replicated twice, as shown in main text and in supplemental material.

### Analysis of Mal3-GFP tip tracking in cell-free experiments

To quantify the Mal3-GFP Specificity, a semi-automated custom MATLAB script was used which allowed the user to pick the position for the peak Mal3 intensity and a position behind the Mal3 comet. To determine the EB1-GFP Specificity (Eqn. 1), a custom MATLAB script was written that allowed the user to pick the brightest point of the comet and then a point on the microtubule lattice, behind the comet. Then, a 4 X 4 pixel box was summed at the brightest point of the comet (*Tip_intensity*) and a separate 4 X 4 pixel box was summed at the point behind the comet (*Lattice_intensity*). Finally, Equation 1 was used to calculate the EB1 Specificity (*S*). Only the brightest comet per growth event was measured and every growth event in a field of view was analyzed. This was done over three replicates and pooled together for each condition.

For the growth rate, kymographs were made from representative growth events. Then, the growth rate was calculated from the kymograph using ImageJ by determining the length of the microtubule at the start and end of each growth event and dividing by the time required to reach the end of that growth event.

### Simulation prediction for fold change in protofilament-edge on-rate caused by Eribulin

To estimate the fold change in protofilament-edge on-rates caused by Eribulin, the protofilament-edge on-rate in the simulation was slowly reduced by 1.41 (√2) fold change over a range from the baseline protofilament-edge on-rate used in the simulation to 1/16 of the baseline protofilament-edge on-rate, gathering the results from at least 30 simulation runs at each condition. Then, simulated videos for each simulation run were generated and semi-automated code used to estimate EB1 Specificity, as described above. Next, the median value of the Specificity for each simulated condition was plotted and compared to the median Mal3- GFP Specificity from the cell-free experiments. Finally, the Specificity from the simulated videos that was closest to the Specificity for each of the experiments (0, 80, and 160 nM eribulin) was used to estimate the relative reduction in protofilament-edge on-rate caused by the presence of Eribulin in the cell free assay. To obtain error bars for the change in protofilament-edge on rate, the cell-free experimental data for each condition (DMSO, 80 nM Eribulin, 160 nM Eribulin) was bootstrapped 300 times and the median data was plotted (Fig S4D). Then, the difference between Q1 and Q3 of this bootstrapped data was used to create the error bars (Fig 4E).

## Acknowledgements

The Gardner laboratory is supported by a National Institutes of Health grant NIGMS R35- GM126974. SJG is supported by the National Institute of Health Training Program T32GM140936. Parts of this work were carried out in the Characterization Facility, University of Minnesota, which receives partial support from the NSF through the MRSEC (Award Number DMR-2011401) and the NNCI (Award Number ECCS-2025124) programs. We thank members of the Gardner, Courtemanche, and Titus laboratories for helpful discussions. We thank Dr. Thomas Surrey for the generous gift of Mal3 constructs. Finally, we thank Eisai Inc. (Cambridge, MA), and in particular Dr. Bruce A. Littlefield, for the gift of Eribulin to contribute to our research effort.

## Data Availability

All data generated or analyzed during this study are included in this manuscript and supporting files. MATLAB scripts are available via the Gardner lab website https://cbs.umn.edu/gardner<u>-</u> <u>lab/software-links</u>.

## Supplemental Figure

**Figure S1:**
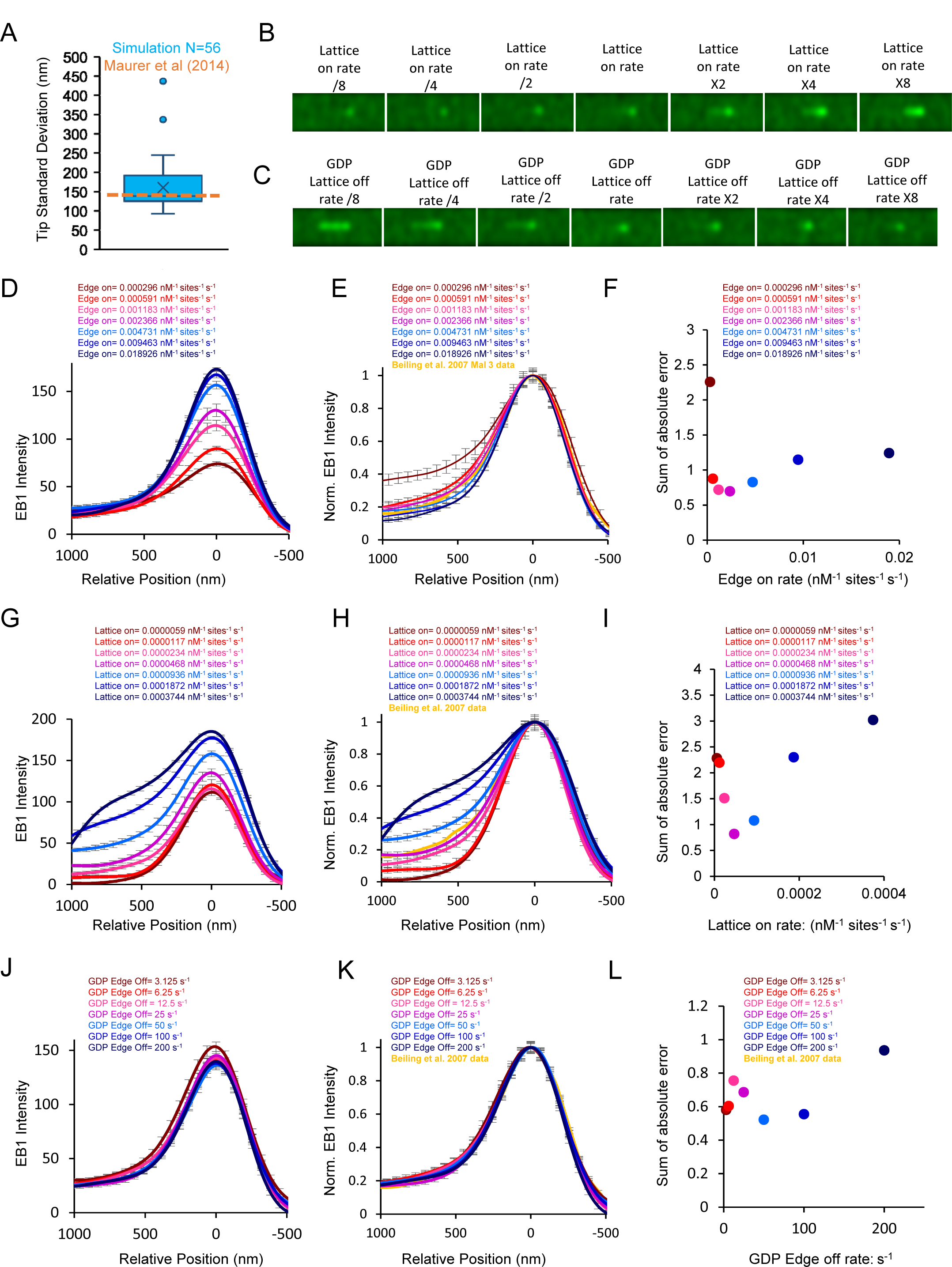
Parameter sensitivity testing for the EB1 tip tracking stochastic model I. (A) Simulated tip standard deviation, a proxy for the variability in protofilament lengths (tip tapering), for the 60 simulated microtubule images that were used to determine EB1 peak position in Fig. 1C. The orange dotted line corresponds to the previously reported experimental fitted tip standard deviation (≤145 nm) (Maurer et al., 2014). (B) Simulated EB1 comets with varying closed-lattice on rates. (C) Simulated EB1 comets with varying GDP closed-lattice off rates. (D) Line scans of simulated EB1 tip tracking with varying protofilament-edge on-rates. (E) Normalized line scans of simulated EB1 tip tracking with varying protofilament-edge on-rates compared to literature data (Bieling et al., 2007). (F) Sum of absolute difference between EB1 line scan for each protofilament-edge on-rate and the literature data. (G) Line scans of simulated EB1 tip tracking with varying closed-lattice on-rates. (H) Normalized line scans of simulated EB1 tip tracking with varying closed-lattice on-rates compared to literature data (Bieling et al., 2007). (I) Sum of absolute difference between EB1 line scan for each closed- lattice on-rate and the literature data. (J) Line scans of simulated EB1 tip tracking with varying GDP protofilament-edge off-rates. (K) Normalized line scans of simulated EB1 tip tracking with varying GDP protofilament-edge off-rates compared to literature data (Bieling et al., 2007). (L) Sum of absolute difference between EB1 line scan for each GDP protofilament-edge off-rate and the literature data.

**Figure S2.**
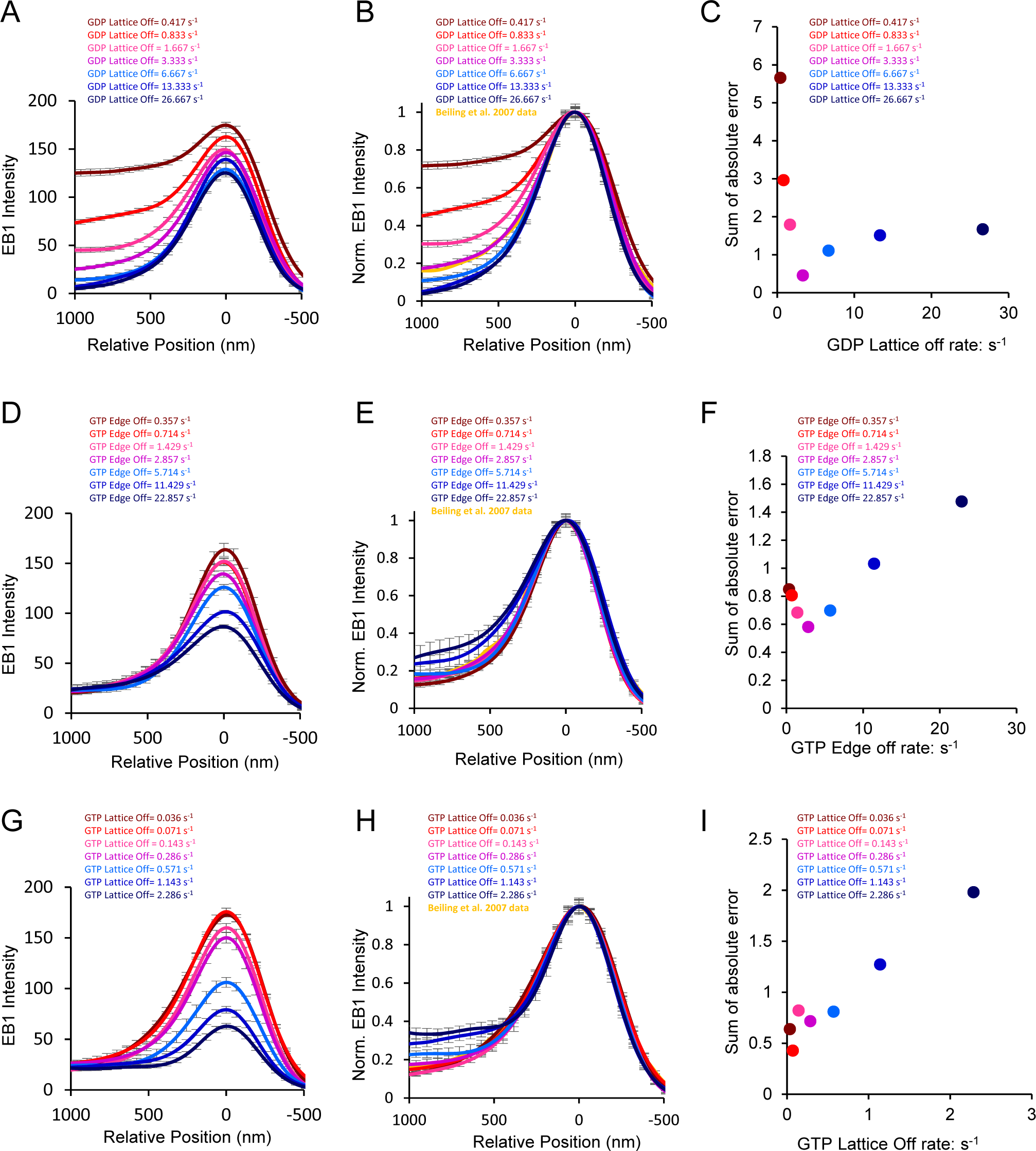
Parameter sensitivity testing for the EB1 tip tracking stochastic model II. (A) Line scans of simulated EB1 tip tracking with varying GDP closed-lattice off-rates. (B) Normalized line scans of simulated EB1 tip tracking with varying GDP closed-lattice off-rates compared to literature data (Bieling et al., 2007). (C) Sum of absolute difference between EB1 line scan for each GDP closed-lattice off-rate and the literature data. (D) Line scans of simulated EB1 tip tracking with varying GTP protofilament-edge off-rates. (E) Normalized line scans of simulated EB1 tip tracking with varying GTP protofilament-edge off-rates compared to literature data (Bieling et al., 2007). (F) Sum of absolute difference between EB1 line scan for each GTP protofilament-edge off-rate and the literature data. (G) Line scans of simulated EB1 tip tracking with varying GTP closed-lattice off-rates. (H) Normalized line scans of simulated EB1 tip tracking with varying GTP closed-lattice off-rates compared to literature data (Bieling et al., 2007). (I) Sum of absolute difference between EB1 line scan for each GTP closed-lattice off-rate and the literature data.

**Figure S3:**
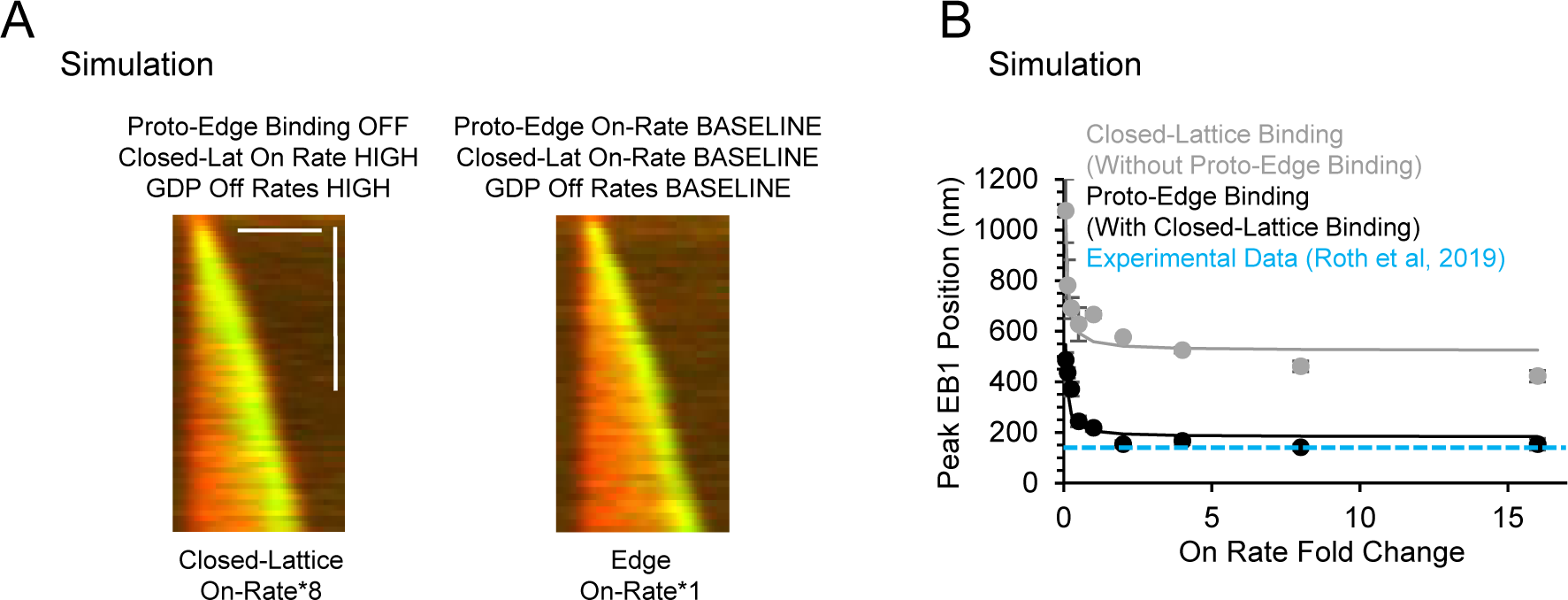
Simulation data for effect of high GDP closed-lattice off-rates. (A) (Left) To determine if closed-lattice binding alone could recapitulate tip tracking, a simulated kymograph was generated with a high closed-lattice on rate, as well as very high GDP-tubulin off-rates. Even in this case, the EB1 comet does not properly localize at the growing microtubule end, as is observed when comparing to the (right) simulated kymograph with the base protofilament edge on rate. (B) The peak EB1 position without protofilament-edge binding ranges from 428- 666 nm distal of the growing microtubule end (grey) compared to the literature value of ∼144 nm (blue dotted line). Meanwhile, the peak EB1 position with protofilament-edge binding ranges from 141-217 nm distal of the growing microtubule end (black), similar to the literature value of ∼144 nm (blue dotted line).

**Figure S4:**
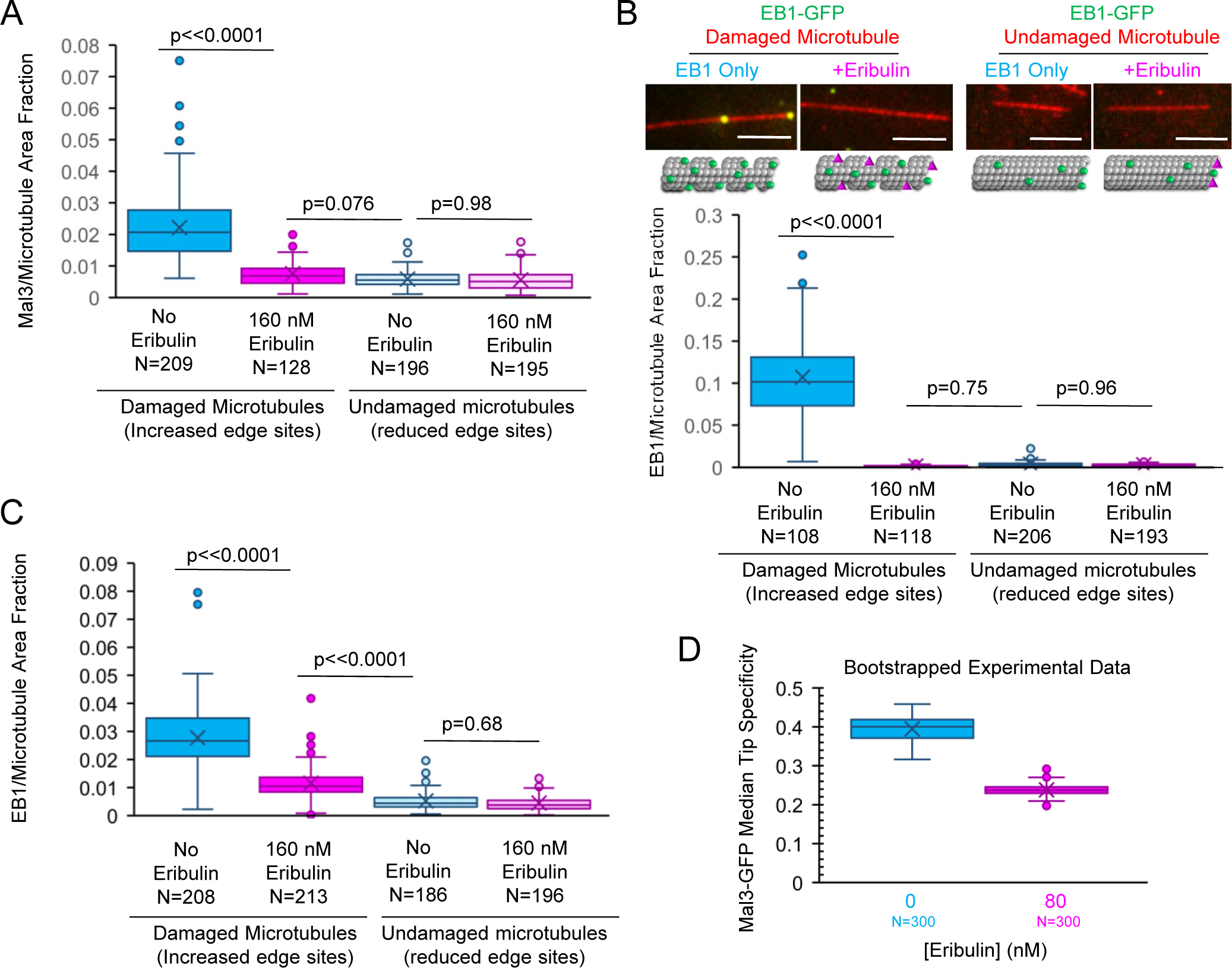
Experimental data for binding of EB1-GFP and Mal3-GFP to stabilized GMPCPP microtubules. A) Quantification of the fraction of microtubule area bound by Mal3-GFP for damaged and undamaged microtubules treated with Mal3-GFP in the absence or presence of Eribulin (ANOVA, p<<0.0001; Tukey’s post-hoc for individual comparisons). (B) Top: EB1-GFP binding to damaged and undamaged GMPCPP microtubules in the absence or presence of Eribulin. 5 μm scale bar. Bottom: Quantification of the fraction of microtubule area bound by EB1-GFP for damaged and undamaged microtubules treated with EB1-GFP in the absence or presence of Eribulin (ANOVA, p<<0.0001; Tukey’s post-hoc for individual comparisons). (C) Quantification of the fraction of microtubule area bound by EB1-GFP for damaged and undamaged microtubules treated with EB1-GFP in the absence or presence of Eribulin (ANOVA, p<<0.0001; Tukey’s post-hoc for individual comparisons). (D) Bootstrapped Mal3 specificity data in the absence and presence of Eribulin.

**Figure S5.**
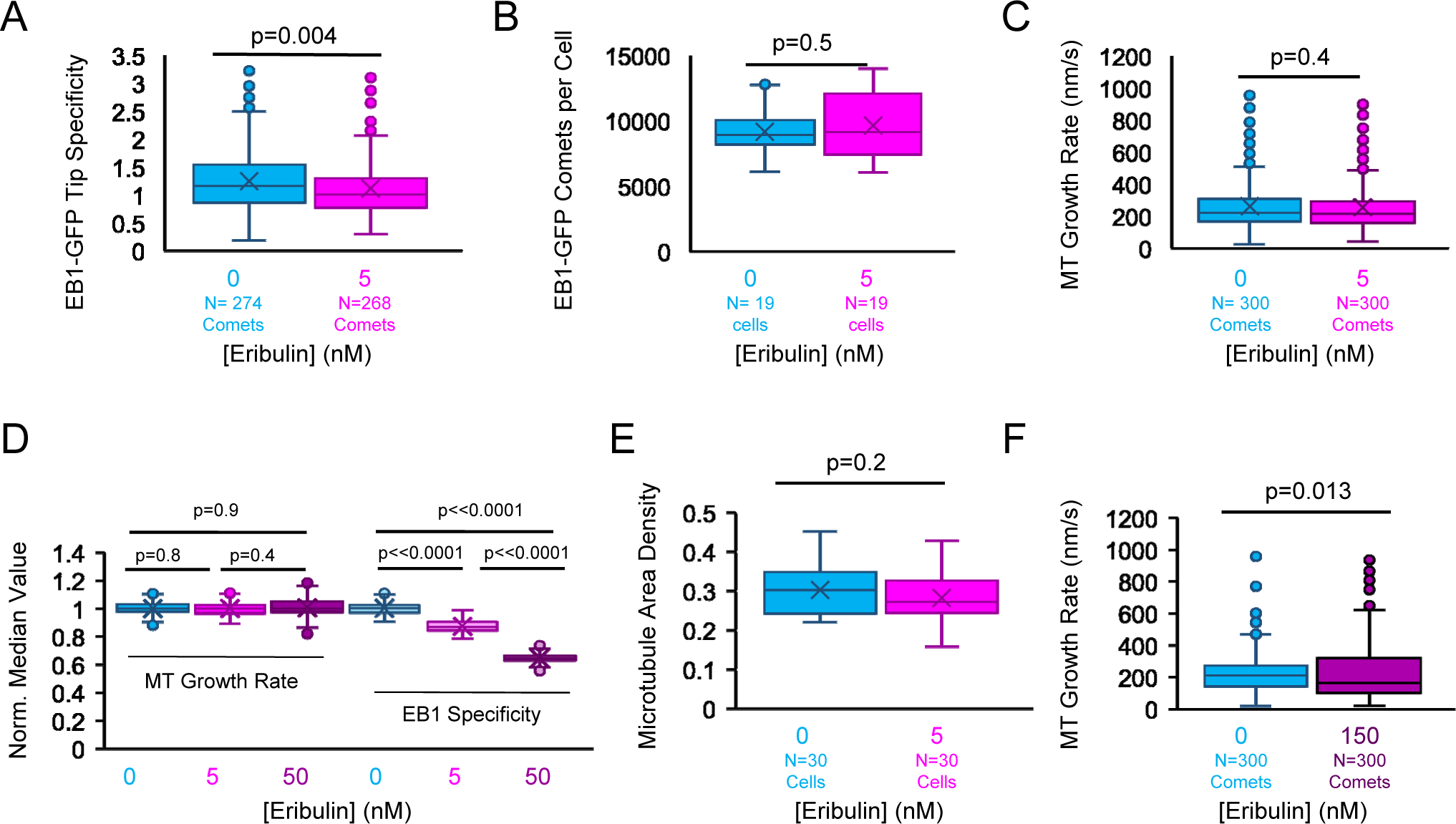
Experimental data in cells, effect of Eribulin on EB1 tip tracking. (A) EB1-GFP Tip Specificity for individual comets in cells treated with 0 or 5 nM Eribulin (t-Test, p<<0.0001). (B) Average number of EB1-GFP comets per cell treated with 0 or 5 nM Eribulin determined using uTrack from the Danuser lab (Applegate et al., 2011) (t-test, p=0.0003). (C) Growth rate of individual comets in cells treated with 0 or 5 nM Eribulin (Mann-Whitney U test, p<<0.0001). (D) Bootstrapped Experimental data to observe the variation in median EB1-GFP specificity and growth rate in cells with varying Eribulin concentrations (ANOVA, p<<0.0001, Tukey’s HSD for other comparisons). (E) Quantification of microtubule density in cells with 0 or 5 nM Eribulin (T-test, p=0.220). (F) Growth rate of individual comets in cells treated with 0 or 150 nM Eribulin (Mann-Whitney U test, p=0.0126).

## Movie Legends

**Movie S1**: Simulated EB1 tip tracking. EB1-GFP in green, Microtubule in red. 2 µm scale bar.

**Movie S2**: Simulated EB1 tip tracking from simulation with different hydrolysis rates. EB1-GFP in green, microtubule in red. 2 µm scale bar. Fast Hydrolysis rate on left (2 s^-1^), slow hydrolysis rate on right (0.1 s^-1^).

**Movie S3**: Simulated EB1 tip tracking over a range of EB1 closed-lattice on rates (1.2x10^-5^ – 3.7x10^-4^ nM^-1^ sites ^-1^ s^-1^) and EB1 protofilament-edge binding on-rate set to 0. For these simulations, EB1 on and off rates were kept constant except for the closed-lattice on-rate, which increases from left to right for each subset of the panel. EB1-GFP in green, microtubule in red. 2 µm scale bars.

**Movie S4**: Simulated EB1 tip tracking over a range of protofilament-edge on-rates (3x10^-4^ – 9.5x10^-3^ nM^-1^ sites^-1^ s^-1^). All EB1 on-and off-rates were kept constant except for the protofilament-edge on-rate, which increases from left to right for each subset of the panel. EB1-GFP in green, microtubule in red. 2 µm scale bars.

**Movie S5**: Simulated EB1 tip tracking, with EB1 closed-lattice binding dynamically set to 0 during the run. Simulation is run with all standard parameters. At time 9 sec, EB1 closed-lattice on-rate is set to zero. Then, at time 23 sec, the EB1 closed-lattice on-rate is reset to its baseline value of 4.7x10^-4^ nM^-1^ sites^-1^ s^-1^. EB1-GFP in green, microtubule in red. 2 µm scale bar.

**Movie S6**: Simulated EB1 tip tracking, with EB1 protofilament-edge binding dynamically set to 0 during the run. Simulation is run with all standard parameters. At time 9 sec, EB1 protofilament-edge on-rate is set to zero. Then, at time 22 sec, the EB1 protofilament-edge on-rate is reset to its baseline value of 2.4x10^-3^ nM^-1^ sites^-1^ s^-1^. EB1-GFP in green, microtubule in red. 2 µm scale bar.

**Movie S7**: Experimental EB1-GFP tip tracking in live LLC-PK1 cells. On left, LLC-Pk1 cells treated with DMSO only, and on right, LLC-Pk1 cells treated with 5 nM Eribulin. EB1-GFP in green. 10 µm scale bar.

**Movie S8**: Experimental EB1-GFP tip tracking in live LLC-PK1 cells. On left, LLC-Pk1 cells treated with DMSO only, and on right, LLC-Pk1 cells treated with 50 nM Eribulin. EB1-GFP in green. 10 µm scale bar.

**Movie S9**: Experimental EB1 tip tracking in live LLC-PK1 cells. On left, LLC-PK1 cells treated with DMSO only, and on right, LLC-PK1 cells treated with 150 nM Eribulin. EB1-GFP in green. 10 µm scale bar.

